# Distinguishing cells from empty droplets in droplet-based single-cell RNA sequencing data

**DOI:** 10.1101/234872

**Authors:** Aaron T. L. Lun, Samantha Riesenfeld, Tallulah Andrews, The Phuong Dao, Tomas Gomes, participants in the 1st Human Cell Atlas Jamboree, John C. Marioni

## Abstract

Droplet-based single-cell RNA sequencing protocols have dramatically increased the throughput and efficiency of single-cell transcriptomics studies. A key computational challenge when processing these data is to distinguish libraries for real cells from empty droplets. Existing methods for cell calling set a minimum threshold on the total unique molecular identifier (UMI) count for each library, which indiscriminately discards cell libraries with low UMI counts. Here, we describe a new statistical method for calling cells from droplet-based data, based on detecting significant deviations from the expression profile of the ambient solution. Using simulations, we demonstrate that our method has greater power than existing approaches for detecting cell libraries with low UMI counts, while controlling the false discovery rate among detected cells. We also apply our method to real data, where we show that the use of our method results in the retention of distinct cell types that would otherwise have been discarded.

## Introduction

Recent advances in droplet-based protocols have revolutionized the field of single-cell transcriptomics by allowing tens of thousands of cells to be profiled in a single assay [1–3]. In these technologies, individual cells are captured into aqueous droplets in a water-in-oil emulsion. Each droplet also contains a co-captured bead with primers for reverse transcription, where all primers on a single bead contain a cell barcode that is (effectively) unique to that bead. The droplets serve as isolated reaction chambers in which cell lysis and reverse transcription are performed to obtain barcoded cDNA. This is followed by breaking of the emulsion, amplification of the cDNA and construction of a sequencing library. After sequencing, transcripts are assigned to individual droplets based on the cell barcode observed in each read sequence. This yields an expression profile for each cell, typically in the form of unique molecular identifier (UMI) counts [4] for all annotated genes. The use of droplets increases throughput by at least an order of magnitude compared to protocols based on plates [5] or conventional microfluidics [6], which is appealing for large-scale projects such as the Human Cell Atlas [7].

The complexity of the sequencing data from droplet-based technologies poses a number of interesting challenges for low-level data processing. One such challenge is the identification and removal of cell barcodes corresponding to empty droplets. An empty droplet does not contain a cell but will still contain “ambient” RNA [1], i.e., cell-free transcripts in the solution in which the cells are suspended. Ambient RNA can be actively secreted by cells or released upon cell lysis (possibly induced by the stresses of dissociation and microfluidics). The presence of ambient RNA means that many empty droplets will contain material for reverse transcription and library preparation, resulting in non-zero total UMI counts for the corresponding barcodes. However, the expression profiles for these barcodes do not originate from any individual cell and need to be removed prior to further analysis to avoid misleading biological conclusions.

Existing methods for removing empty droplets assume that droplets containing genuine cells should have more RNA, resulting in larger total UMI counts for the corresponding barcodes. Zheng *et al*. [3] remove all barcodes with total counts below 10% of the 99^th^ percentile of the Y largest total counts, where *Y* is defined as the expected number of cells to be captured in the experiment. Macosko *et al*. [1] set the threshold at the knee point in the cumulative fraction of reads with respect to increasing total count. While simple, the use of a one-dimensional filter on the total UMI count is suboptimal as it discards small cells with low RNA content. Droplets containing small cells are not easily distinguishable from empty droplets based on the total number of transcripts. This is due to variable capture and amplification efficiencies across droplets during library preparation, which mixes the distributions of total counts between empty and non-empty droplets. Applying a simple threshold on the total count forces the researcher to choose between the loss of small cells or an increase in the number of artifactual “cells” composed of ambient RNA. This is especially problematic if small cells represent distinct cell types or functional states.

Here, we propose a new method for detecting empty droplets in droplet-based single-cell RNA sequencing (scRNA-seq) data. We estimate the profile of the ambient RNA pool and test each barcode for deviations from this profile using a Dirichlet-multinomial model of UMI count sampling. Barcodes with significant deviations are considered to be genuine cells, thus allowing recovery of cells with low total RNA content and small total counts. We combine our approach with a knee point filter to ensure that barcodes with large total counts are always retained. Using a variety of simulations, we demonstrate that our method outperforms methods based on a simple threshold on the total UMI count. We also apply our method to several real datasets where we are able to recover more cells from both existing and new cell types.

## Description of the method

### Testing for deviations from the ambient profile

To construct the profile for the ambient RNA pool, we consider a threshold *T* on the total UMI count. The set 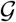 of all barcodes with total counts less than or equal to *T* are considered to represent empty droplets. The exact choice of *T* does not matter, as long as (i) it is small enough so that droplets with genuine cells do not have total counts below *T*, and (ii) there are sufficient counts to obtain a precise estimate of the ambient profile. We set *T* = 100 by default in our approach, motivated by examination of several real datasets (Supplementary Section 1, Supplementary Figure 1). We stress that *T* is not the same as the threshold used in existing methods, as barcodes with total counts greater than *T* are not automatically considered to be cell-containing droplets.

The ambient profile is constructed by summing counts for each gene across 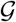. Let *y_gb_* be the count for gene *g* in barcode *b*. We define the ambient count for *g* as

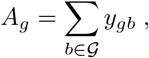

yielding a count vector **A** = (*A*_1_,…, *A_N_*) for all *N* genes. (We assume that any gene with counts of zero for all barcodes has already been filtered out, as this provides no information for distiguishing between barcodes.) We apply the Good-Turing algorithm to **A** to obtain the posterior expectation, 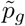, of the proportion of counts assigned to *g* [8], using the `goodTuringProportions` function in the `edgeR` package [9]. This ensures that genes with zero counts in the ambient pool have non-zero proportions, avoiding likelihoods of zero in downstream calculations. In general, we do not observe strong differential expression between **A** and the average of the cell-containing droplets (Supplementary Figure 2). This suggests that the ambient pool contains RNA from multiple cell types, possibly from widespread stress and lysis during dissociation.

Our null hypothesis is that free-floating transcripts in solution are randomly encapsulated into the empty droplets. For a given droplet, the probability of sampling a transcript molecule for gene *g* is equal to 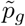. If we condition on the total count *t_b_* for a cell barcode *b*, we can model the counts for each barcode with a Dirichlet-multinomial distribution. (A physical justification for this model is provided in Supplementary Section 2.) We define the likelihood of obtaining the counts for barcode *b* as

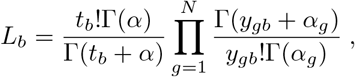

where 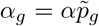 for a scaling factor *α*. A maximum likelihood estimate of *α* is obtained from 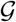 by treating 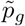 as known. Low estimates of *α* model overdispersion in the counts, e.g., due to amplification biases or correlated sampling of transcript molecules.

We use a Monte Carlo approach to compute the *p*-value for *b*. At each iteration *i*, we generate a new vector of counts by randomly sampling from a Dirichlet-multinomial distribution with probabilities set to 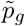 for all *g*, the size set to *t_b_*, and the scaling factor set to (our estimate of) *α*. We calculate the likelihood 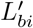 for this count vector using the same expression as shown above for *L_b_*. We repeat this process for *R* iterations and then use the method of Phipson and Smyth [10] to define the *p*-value as

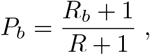

where *R_b_* is the number of iterations in which 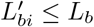. This strategy avoids *p*-values of zero, which is important during multiple testing correction. See Supplementary Section 3 for a description of how these *p*-values are efficiently computed.

### Detecting the knee point in the log-totals

Applying a threshold on the *p*-value will identify barcodes that have count profiles that are significantly different from the ambient pool of RNA. We assume that this will be the case for most cell-containing droplets, as the ambient pool is formed from many (lysed) cells and is unlikely to be representative of any single cell. However, it is possible for some cell-containing droplets to have ambient-like expression profiles. This can occur if the cell population is highly homogeneous or if one cell subpopulation contributes disproportionately to the ambient pool, e.g., if it is more prone to lysis. Sequencing errors in the cell barcodes may also bias the estimates of the ambient proportions, by misassigning counts from cell-containing droplets to barcodes with low UMI totals. This may result in spurious similarities between cells and the estimated ambient profile.

To avoid incorrectly calling ambient-like cells as empty droplets, we combine our procedure with a conventional threshold on the total UMI count. We rank all barcodes in order of decreasing *t_b_*, and consider log(*t_b_*) as a function *f*(.) of the log-transformed rank, i.e., log(*t_b_*) = *f*(log *r_b_*) where *r_b_* is the rank of *b* in the ordered sequence of barcodes. The first “knee” point in this function corresponds to a transition between a distinct subset of barcodes with large totals and the majority of barcodes with smaller totals. This is defined at the log-rank that minimizes the signed curvature

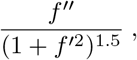

and represents the point at which *f*(.) begins to drop rapidly, marking the start of the transition between large and small totals. In practice, we obtain *f*(.) by fitting a smooth spline to log(*t_b_*) against the log-rank in the interval containing the knee point. The derivatives of *f*(.) are then obtained by differentiation of the spline basis functions. This avoids multiplication of errors during numerical differentiation, which would lead to instability in the curvature values and inaccurate estimates of the knee point.

Our assumption is that any barcode with a large total count must represent a cell-containing droplet, regardless of whether its count profile resembles the ambient pool. This is based on the expectation that the distribution of the sizes of empty droplets should be unimodal, with a monotonic decreasing probability density as *t_b_* increases past the mode. A distinct peak of large totals would not be consistent with this expected distribution. We define the upper threshold *U* as the *t_b_* at the knee point and retain all barcodes with *t_b_* ≥ *U*, irrespective of their *P_b_*. This guarantees recovery of any barcodes with large total counts that potentially represent cell-containing droplets. We use the knee point rather than the inflection point as the *t_b_* at the former is larger, providing a more conservative threshold that avoids retention of empty droplets.

We stress that, despite the use of a threshold on *t_b_*, our approach is different from existing methods due to the testing procedure. Barcodes with *t_b_* below the knee point can still be retained if the count profile is significantly different from the ambient pool. This is not possible with existing methods that would simply discard these barcodes. Users can also set *U* manually if automatic detection of the knee point fails for complex *f*(.). Alternatively, this mechanism can be disabled completely in favour of detecting cells solely based on their *p*-values. This is more statistically rigorous as it avoids the selection of an *ad hoc* threshold, but may result in the failure to detect large cells.

### Correcting for multiple testing across barcodes

We correct for multiple testing by controlling the false discovery rate (FDR) using the Benjamini-Hochberg (BH) method [11]. Putative cells are defined as those barcodes that have significantly poor fits to the ambient model at a specified FDR threshold. In the following text, we will use an FDR threshold of 0.1% unless otherwise mentioned. This means that the expected proportion of empty droplets in the set of retained barcodes is no greater than 0.1%, which we consider to be acceptably low for downstream analyses. Users are also free to choose their own thresholds, with more relaxed thresholds favouring sensitivity in cell detection at the cost of specificity.

Note that we only perform the BH correction on the *p*-values for barcodes that have *t_b_* greater than *T*. This reduces the severity of the correction by discarding barcodes that were previously assumed to be empty droplets, thus improving detection power for barcodes with larger totals that are more likely to contain cells. In fact, *p*-values are not computed at all for barcodes with *t_b_* ≤ *T* to avoid unnecessary computational work. Conversely, all barcodes with *t_b_* ≥ *U* are considered to be known true positives, regardless of how ambient-like their expression profiles are. These barcodes have their *p*-values set to zero during the BH correction. This approach improves power by reducing the severity of the correction in the presence of a set of known positives.

## Results

### Evaluating performance with simulated droplet-based data

We named our method “EmptyDrops” and tested it on simulated data involving cells with different RNA content (see Methods, Supplementary Figure 3). Each simulated dataset was generated from real droplet-based scRNA-seq data (Supplementary Table 2) and contained one group of large cells with high RNA content and large *t_b_*; one group of small cells with low RNA content and small *t_b_*; and a set of empty droplets with counts sampled from an ambient pool of RNA. We applied EmptyDrops at an FDR of 0.1% to determine the recall for each group of cells and the FDR among the detected barcodes. We also tested methods that retain all cells with total UMI counts above a threshold. The threshold was defined as the total *U* at the knee point, as described above; or using the quantile-based approach [3] in the CellRanger software from 10X Genomics.

In a simulation based on a real dataset containing peripheral blood mononuclear cells (PBMCs), EmptyDrops detected the most cells from both groups (Figure 1). CellRanger and the knee point method detected large cells but failed to recover small cells. We observed similar results in simulations based on other real datasets (Supplementary Figures 4-8). The poor performance of the total count-based methods for small cells is expected. Barcodes corresponding to small cells with little RNA have similar total counts as barcodes corresponding to empty droplets with many ambient molecules. A method based on the total count alone cannot distinguish between these two possibilities, as any choice of threshold will either reduce recall or increase false positives (Supplementary Figure 9). In contrast, EmptyDrops uses the expression profile for each droplet to distinguish small cells from the ambient profile with greater power.

**Figure 1.**
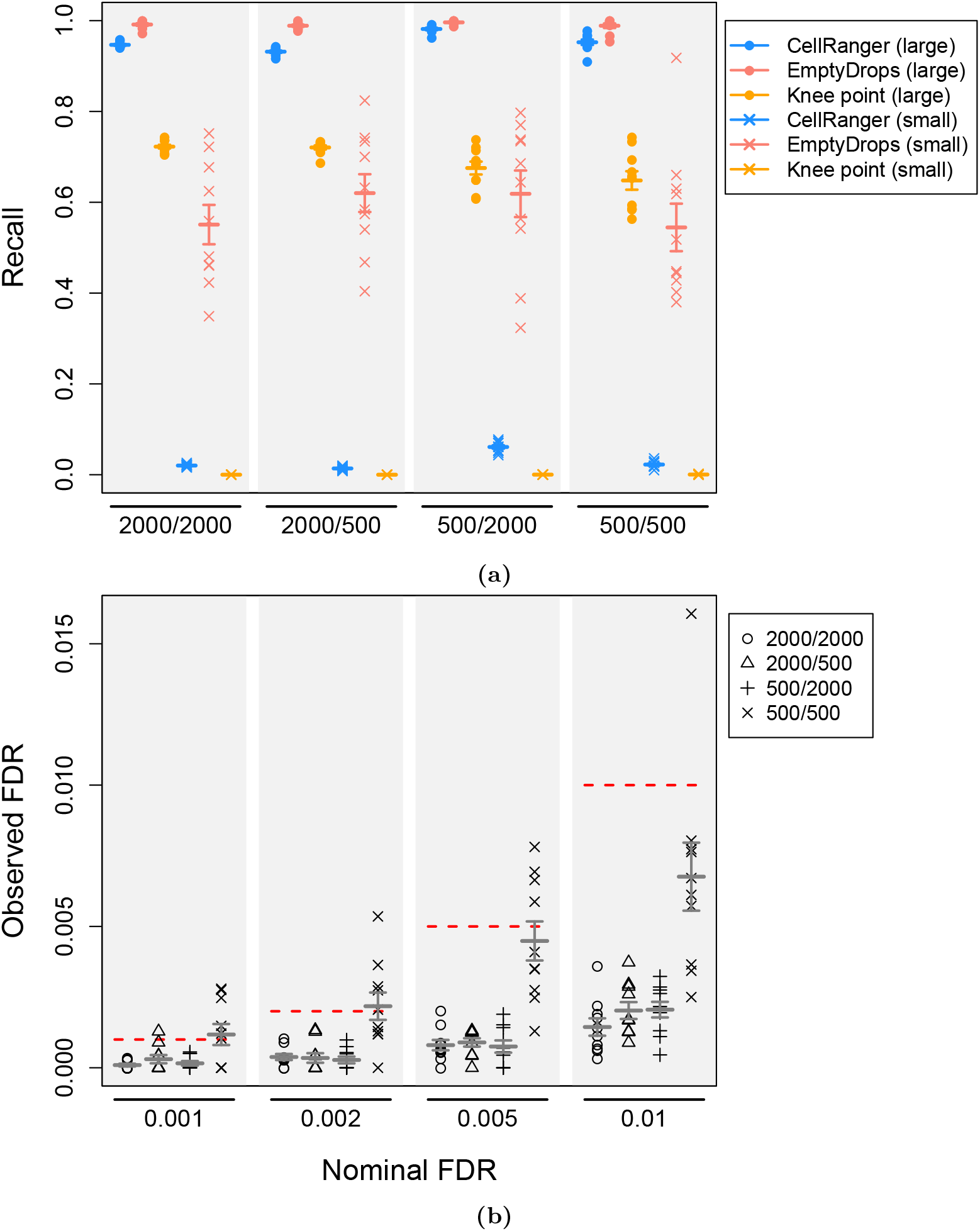
Cell-calling results from different algorithms in simulations based on the PBMC dataset. Simulation scenarios are labelled as *G*_1_/*G*_2_ where *G*_1_ and *G*_2_ are the number of barcodes in the group of large and small cells, respectively. (a) The recall for each method, defined as the proportion of detected cells from each group. EmptyDrops was used with an FDR threshold of 0.1%. (b) The observed FDR in the set of libraries detected by EmptyDrops at a range of nominal FDR thresholds (dotted lines), defined as the proportion of detected droplets that are empty. In both plots, each point represents the result of one simulation iteration, the bar represents the mean value across 10 iterations and the error bars represent the standard error of the mean.

EmptyDrops correctly controlled the FDR close to or below the nominal thresholds in all simulated datasets (Figure 1, Supplementary Figures 4-8). This is a useful property of the method as it provides users with a reliable upper bound on the expected proportion of empty droplets. Such information can be used to interpret downstream analysis results – for example, we would be satistifed that a clustering result was not driven by empty droplets if the proportion of cells in a cluster of interest was much higher than the FDR threshold used in EmptyDrops. By comparison, the effect of a total count threshold on the cell calling error rate is less obvious. CellRanger also requires the expected number of cells, which may not be available or accurate.

### Characterizing behaviour of EmptyDrops on real datasets

To determine how EmptyDrops behaved on real data, we applied it to detect cells in a placenta dataset [12] at an FDR of 0.1% (Figure 2). EmptyDrops identified a visually appropriate *U* using the knee point from the smoothed spline (Figure 2a), and detected significant barcodes as those with low likelihoods under the null Dirichlet-multinomial model (Figure 2b). Most of the barcodes detected as cells by EmptyDrops had large total counts and were also detected using CellRanger (Figure 2c). Barcodes that were only detected by EmptyDrops had low total counts (Figure 2d), consistent with the expected differences between methods. We observed similar results in the other tested datasets (Supplementary Table 2, Supplementary Figures 10-15) where EmptyDrops often detected the most barcodes. Increased retention of small cells was particularly pronounced in the neuronal datasets where EmptyDrops uniquely detected over a thousand cells. A smaller number of barcodes were uniquely detected by CellRanger in a few datasets, the causes of which are discussed in Supplementary Section 4.

**Figure 2.**
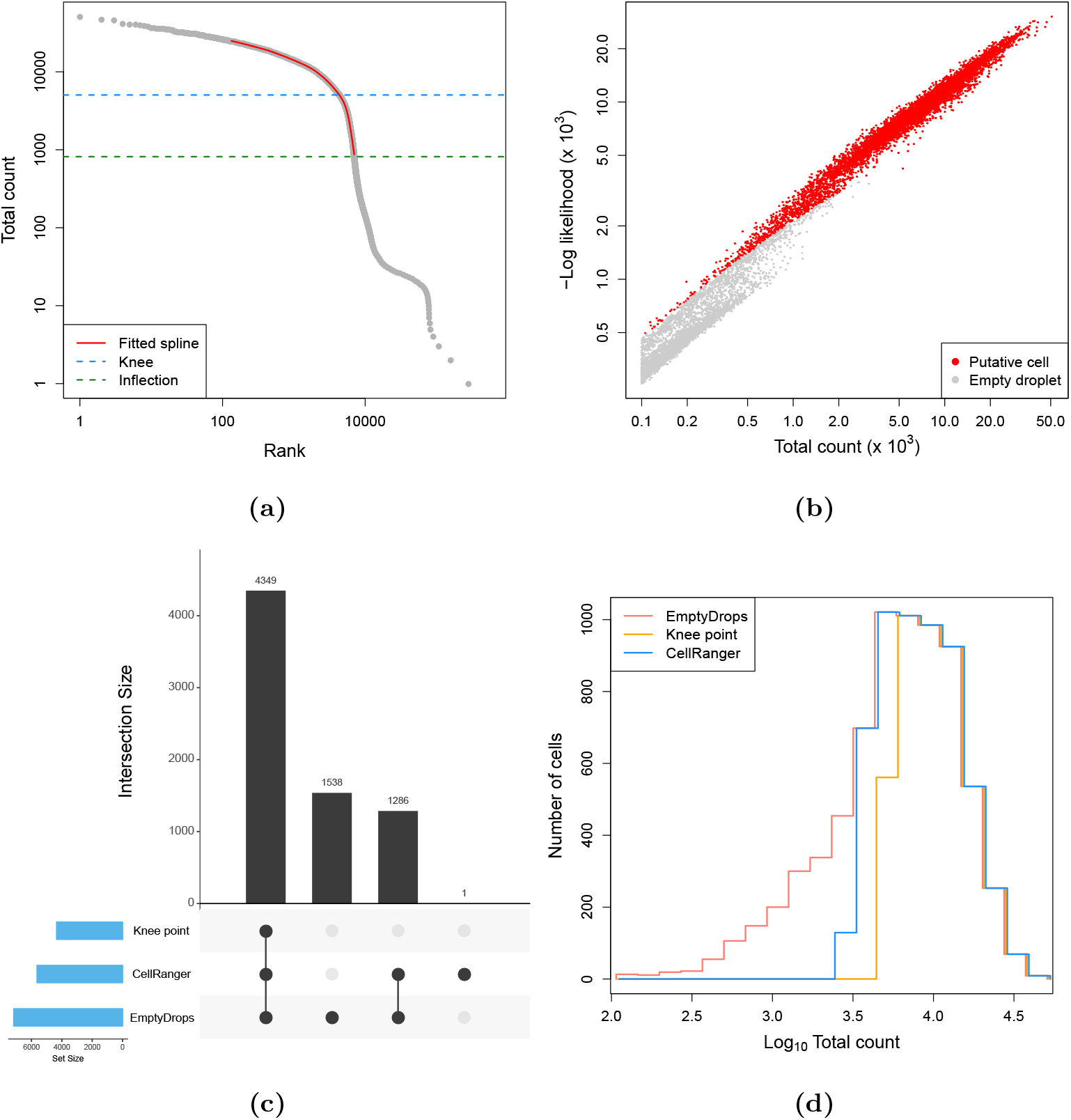
Application of EmptyDrops and other cell detection methods to one sample of the placenta dataset. (a) A barcode rank plot showing the fitted spline used for knee point detection in EmptyDrops. The detected knee and inflection points are also shown. (b) The negative log-likelihood for each barcode in the multinomial model of EmptyDrops, plotted against the total count. Barcodes detected as putative cell-containing droplets at an FDR of 0.1% are labelled in red. Only barcodes with *t_b_* > *T* are shown. (c) An UpSet plot [13] of the barcodes detected by each combination of methods (vertical bars). Horizontal bars represent the number of barcodes detected by each method. (d) Histogram outlines of the log-total count for barcodes detected by each method.

To explore the differences between methods in more detail, we generated *t*-stochastic neighbour embedding (t-SNE) plots [14] of all barcodes that were detected by either CellRanger or EmptyDrops in several datasets. In the placenta dataset, many of the EmptyDrops-only barcodes formed unique clusters (Figure 3a), one of which likely contains monocytes (Figure 3b). This suggests that the use of EmptyDrops enables the recovery of distinct cell types, which is not surprising as the total RNA content of a cell is often associated with its biology. We repeated our analysis on another 10X dataset containing approximately 900 brain cells. EmptyDrops uniquely retained a large number of barcodes, including a putative cluster of interneurons (Figures 3c, d) that would have been lost with CellRanger. This may reflect the difficulty of dissociating brain tissue without loss of cytoplasmic RNA [15] that yields low total counts in the resulting libraries. In the PBMC dataset, the EmptyDrops-only barcodes again formed a separate cluster corresponding to platelet-like cells (Supplementary Figure 16). This is consistent with the fact that platelets have much less RNA than other cell types [16].

**Figure 3.**
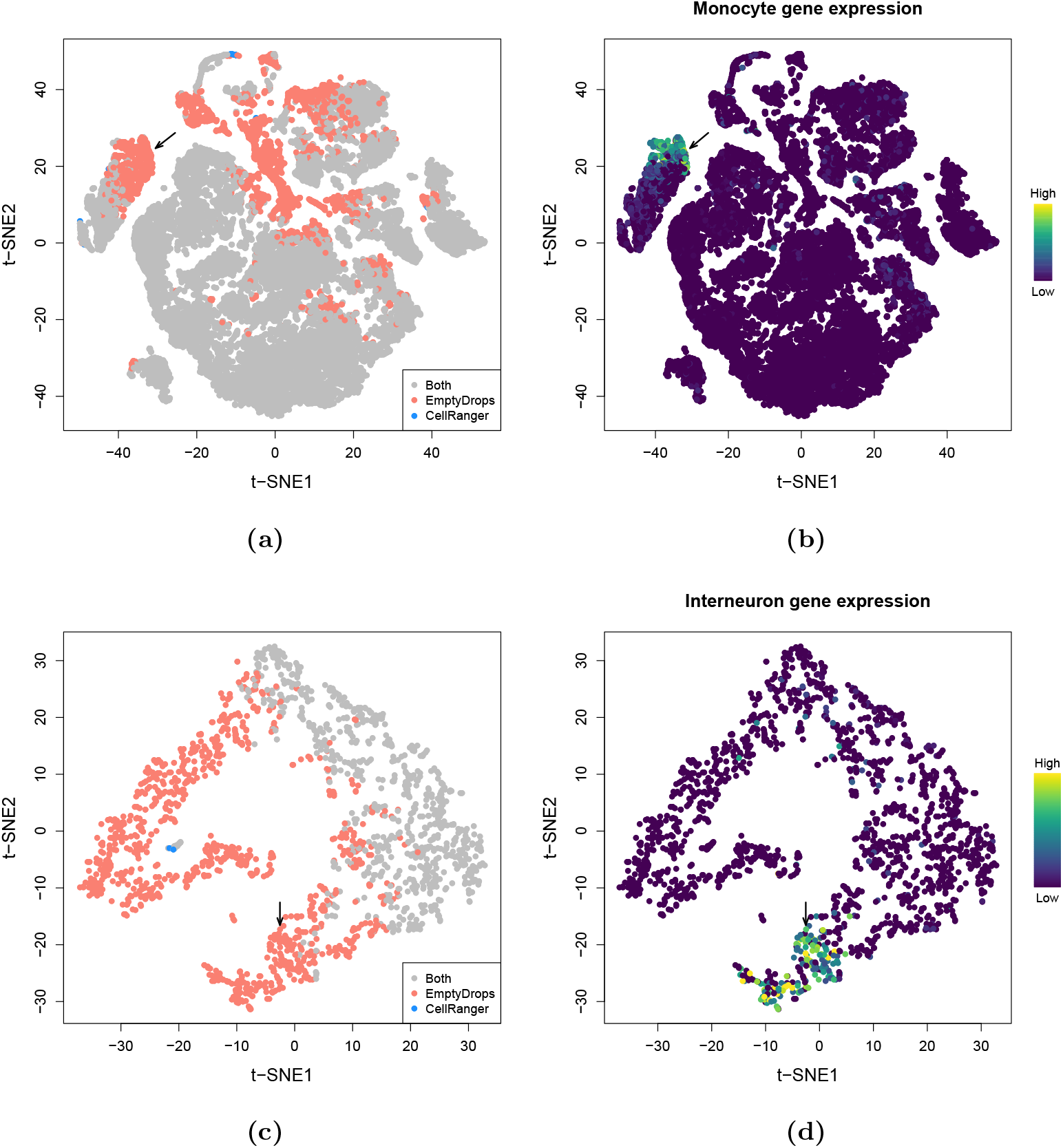
*t*-SNE plots for the placenta dataset (a, b) or the 900 neuron dataset (c, d), constructed from barcodes that were detected with EmptyDrops and/or CellRanger. Each point represents a barcode and is coloured based on (a, c) whether it was detected as a cell with each method; (b) the expression of monocyte marker genes *KCNA5, CFP, STX11* and *S100A12*; or (d) the expression of interneuron marker genes *Gad1, Gad2* and *S1a6c1*. Expression of the relevant marker set in each barcode was quantified as the sum of the normalized log-expression values across all marker genes. Arrows mark the putative monocyte and interneuron populations in each dataset.

## Discussion

Droplet-based technologies are becoming increasingly popular for high-throughput single-cell transcriptomics. However, little work has been performed to develop robust computational methods for distinguishing genuine cells from empty droplets. Here, we describe EmptyDrops, a method to detect cell-containing barcodes based on significant deviation of the expression profiles from the pool of ambient RNA. We use simulated data to demonstrate that EmptyDrops outperforms the strategy currently implemented in the CellRanger software suite. EmptyDrops can recover biology in real 10X data that is lost using CellRanger. Our results indicate that EmptyDrops is effective for cell detection in droplet-based scRNA-seq data. This is supported by other work where EmptyDrops improves cell type recovery [17] and reduces technical artifacts [18].

A key assumption of our approach is that barcodes with very low UMI totals represent empty droplets. This allows us to use these barcodes to estimate the ambient profile. However, this assumption may not be appropriate if the dataset contains a subset of cells with very low RNA content. In such cases, the estimate of the ambient expression profile will be biased, though this bias is likely to be small as few transcripts will be contributed from cells with low RNA content. Another potential source of bias may arise from sequencing errors in the cell barcode, such that transcripts from a cell-containing droplet are misassigned to an empty droplet. This effect is mitigated by the use of designed cell barcodes in the GemCode protocol, which allows for error correction based on a “whitelist” of known barcode sequences [3]. However, it may be a problem in protocols where error correction of the barcodes is not possible [1].

A notable side-effect of retaining barcodes with low UMI totals is that a higher number of low-quality cells are also recovered. EmptyDrops is technically correct in retaining the associated barcodes as damaged cells are distinct from empty droplets. However, these cells are usually not of interest in downstream analyses. We have removed them by thresholding on the mitochondrial content (see Methods), though other metrics could be used such as the proportion of ribosomal protein mRNA (e.g., if damage has stripped the cytoplasm entirely). If this is not sufficient, manual inspection of the clustering results may be necessary to identify these cells and exclude them from further consideration. The other option is to apply a more stringent threshold on the total count, though this will also discard genuine cells with low RNA content and offset the benefits of using EmptyDrops. Even so, EmptyDrops still provides an advantage over existing methods by providing a statistically rigorous framework for cell detection, without requiring any *a priori* knowledge of the expected number of cells.

We have focused exclusively on droplet-based scRNA-seq data generated using the GemCode technology from 10X Genomics. This is motivated by the widespread use of this platform as well as the availability of the unfiltered datasets (see Methods). In principle, the method can also be applied to data from other droplet-based protocols such as inDrop and Drop-seq. Cell lysis or leakage will occur in any protocol involving dissociation and microfluidics, and the formation of empty droplets containing RNA from the ambient pool is unlikely to be a phenomenon that is unique to 10X datasets.

An interesting direction for future work is whether the contribution of the ambient profile can be “subtracted” from each barcode’s expression profile, thus yielding a more accurate representation of each cell’s transcriptome [19]. This requires estimation of the relative amounts of the ambient pool and cellular RNA in each droplet, which is not straightforward as the ambient pool is itself derived from cells. Accurate quantification of the ambient contribution to each droplet requires ambient-specific “markers” that may not be available for an arbitrary dataset. Direct subtraction of the contribution from the counts is also unsatisfactory as it does not preserve the mean-variance relationship or the uncertainty of the ambient estimates. It seems that an identity-link factor model for count data may be required, which is not trivial to implement.

Our EmptyDrops method is implemented in the `DropletUtils` package, available from the Bioconductor project [20]. We anticipate that it will be useful to researchers who want to extract much information as possible from their droplet-based datasets.

## Methods

### Obtaining droplet-based scRNA-seq datasets

The placenta dataset was obtained from the authors [12] and is available at https://jmlab-gitlab.cruk.cam.ac.uk/publications/EmptyDrops2017-DataFiles. All other datasets were downloaded from the 10X Genomics website (https://support.10xgenomics.com/single-cell-gene-expression/datasets).

Supplementary Table 2 contains a brief summary of the 10X-supplied datasets used in our study. In all cases, only the “raw” count matrices were used to ensure that CellRanger filtering was not already applied to the cell barcodes.

### Evaluating performance with simulated data

For a given real dataset, we computed the total sum of UMI counts *t_b_* for each barcode. We identified the inflection point in the curve of log(*t_b_*) against the log-rank using the `barcodeRanks` function from the `DropletUtils` package. The set of all barcodes with log(*t_b_*) below the inflection point was defined as the set of empty droplets 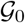. Counts for all 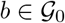 were summed together to create an ambient pool of RNA molecules. (The inflection point provides a conservative definition of empty droplets, and avoids the inclusion of cell-containing droplets in the ambient pool.) To simulate known empty droplets, we constructed expression profiles for a new set of barcodes by sampling molecules from the ambient pool without replacement. This was done such that the distribution of *t_b_* in our set was the same as that in 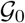. In this manner, we recapitulated the observed number of empty droplets and their total counts in our simulations.

To obtain *G*_1_ large cells, we sampled from the set of barcodes with log(*t_b_*) above the inflection point. We used sampling with replacement to avoid problems in cases where *G*_1_ is greater than the number of estimated cells in the dataset. To generate *G*_2_ small cells, we sampled from the same set of barcodes and downsampled the count vector for each barcode to 10% of its original total, using the `downsampleMatrix` function from the `DropletUtils` package. This mimics the presence of small cells with low RNA content. In both cases, we scrambled a small proportion (10%) of randomly selected genes to eliminate any similarities to the ambient pool in the sampled set of profiles. This ensures that the assumed true cells in our simulation are not actually empty droplets. We tested different simulation scenarios by setting *G*_1_ or *G*_2_ to 500 and 2000 cells. The various components of the simulation are visualized in Supplementary Figure 3.

We applied our EmptyDrops method to the simulated data at an FDR of 0.1%. The recall was defined as the proportion of known cells from each group that were successfully detected. The observed false discovery rate was defined as the proportion of detected barcodes that were known empty droplets. We repeated this evaluation using the knee point approach, where all barcodes with total counts above the knee point were retained; and with the CellRanger approach, implemented as described [3] with the expected number of cells set to *G*_1_ + *G*_2_ (i.e., the true number of simulated cells).

We generated simulated data based on each real dataset in Supplementary Table 2. For each scenario and dataset, we repeated the simulation for 10 iterations. We used each method in each iteration and collected performance metrics across all iterations.

### Detecting cells in real data with different methods

For each real dataset, we applied EmptyDrops to detect cells at an FDR of 0.1%. We also used the CellRanger approach where the expected number of cells was set to the reported value in Supplementary Table 2; and the knee point method, where the threshold on the total count was defined as the detected knee point in the barcode rank plot. `UpSet` plots were created with using the UpSetR package [13].

### Characterizing detected cells in real datasets

We analyzed the placenta dataset by adapting an existing workflow for scRNA-seq data analysis [21]. We performed the analysis on the union of all cells detected by either CellRanger or EmptyDrops to simplify downstream comparisons between the two methods. First, we removed low-quality cells with high proportions of mitochondrial reads by detecting outliers based on the median absolute deviation [22]. We calculated cell-specific size factors using the deconvolution method with pre-clustering [23]. We used the size factors to obtain normalized log-expression values for further analysis.

We calculated the biological contribution of the variance for each gene, assuming Poisson technical noise when modelling the mean-variance trend. We performed principal components analysis on the log-expression matrix using the irlba package. We used the first few components as a low-rank approximation of the matrix to speed up downstream steps. The exact number of components was determined using the `denoisePCA` function in `scran`, which matches the sum of biological contributions across all genes to the variance explained by the chosen number of components.

We clustered cells by creating a shared nearest neighbours graph [24] and detecting communities with the Walktrap algorithm from the `igraph` package. Clusters enriched for EmptyDrops-only cells were characterised by detecting differentially expressed genes against every other cluster, using pairwise *t*-tests in the `findMarkers` function from `scran`. A t-SNE plot [14] was generated using from the `Rtsne` and scater packages [22]. We used a perplexity of 30, though similar plots were obtained with other values.

We performed similar analyses on the PBMC and 900 brain cell datasets.

### Implementation details

EmptyDrops is implemented as the `emptyDrops` function in the `DropletUtils` package, available from version 3.8 of the Bioconductor project (https://bioconductor.org/packages/DropletUtils). It is written in a combination of R and C++ and requires approximately 1-2 minutes to run on each of the tested datasets. All code for simulations and real data analysis were written in R and are available at https://github.com/MarioniLab/EmptyDrops2017.

## Supporting information

## Author contributions

ATLL, SR, TA, TPD and TG developed the initial EmptyDrops algorithm and tested it on simulated data. TPD tested the algorithm on the PBMC dataset and TG tested it on the placenta dataset. ATLL improved the efficiency of the algorithm, incorporated it into a package, prepared new simulations for further testing, refined the real data analysis and wrote the manuscript. JCM provided guidance for the project direction.

## Acknowledgements

We would like to thank Jonathan Griffiths for further testing of the algorithm; Elia Benito-Gutierrez for assistance with identifying neuronal markers; Roser Vento-Tormo and Mirjana Efremova for providing the unprocessed placenta dataset and assisting in its interpretation; and Stephen Sansom for discussions on the nature of cell damage. We would also like to thank the AWS Cloud Credits for Research Program for providing computational resources during the Jamboree.

