## Supplementary material for "Distinguishing cells from empty droplets in droplet-based single-cell RNA sequencing data"

#### Supplementary Materials

by

Aaron T. L. Lun[1,*,†], Samantha Riesenfeld[2,*], Tallulah Andrews[3,*], The Phuong Dao[4,*], Tomas Gomes[3,*], participants in the 1st Human Cell Atlas Jamboree[‡], John C. Marioni[1,3,5,#]

[1]Cancer Research UK Cambridge Institute, University of Cambridge, Li Ka Shing Centre, Robinson Way, Cambridge CB2 0RE, United Kingdom

[2]Klarman Cell Observatory, Broad Institute of MIT and Harvard, Cambridge, Massachusetts, USA

[3]Wellcome Trust Sanger Institute, Wellcome Genome Campus, Hinxton, Cambridge CB10 1SA, United Kingdom

[4]Program for Computational and Systems Biology, Sloan Kettering Institute, Memorial Sloan Kettering Cancer Center, New York, NY

[5]EMBL European Bioinformatics Institute, Wellcome Genome Campus, Hinxton, Cambridge CB10 1SD, United Kingdom

* These authors contributed equally to this work.
†
‡ The full list of participants is provided in Supplementary Table 1.
###

December 4, 2018

### 1 Motivating the choice of the total count threshold $T$

The threshold $T$ should be chosen so that cell-containing droplets are not used to estimate the ambient profile. Otherwise, our estimate will be distorted and we will have less power to discriminate between cells and empty droplets. To check if this occurs in real data, we calculated a $p$-value against the ambient null hypothesis for each barcode $b \in \mathcal{G}$, i.e., with $t_b \leq T$ where $T = 100$ by default. Ideally, we should observe a uniform distribution of $p$-values under the assumption that all barcodes in $\mathcal{G}$ are genuinely empty. However, if cell-containing droplets are present in $\mathcal{G}$, we should observe an enrichment of low $p$-values. These low $p$-values either correspond to the cell-containing droplets themselves, or to genuinely empty droplets that no longer fit to our distorted estimate of the ambient profile.

In most datasets, we observed a minor enrichment of low $p$-values in an otherwise uniform distribution (Supplementary Figure 1). We attribute these small spikes of low $p$-values to the presence of cell-containing droplets that fall below the default $T = 100$. Nonetheless, the distributions are mostly uniform, which demonstrates that our statitical model provides an accurate description of transcript sampling. It also suggests that distortions of the ambient profile due to cell-containing droplets are not a major issue. The exception is the `neurons_900` dataset where we see a clear enrichment of very low and very large $p$-values. This effect is not consistent with distortion of the ambient profile, which should only result in a skew towards low $p$-values. Rather, we hypothesize that it is driven by violations of the independent sampling assumption (e.g., if transcript molecules are bound together in RNA complexes and co-sampled into droplets). Positive correlations between molecules would increase the probability of obtaining droplets that are very similar or very different to the ambient profile.

### 2 Physically justifying the Dirichlet-multinomial model

Denote $\lambda_{bg}$ as the number of transcript molecules of gene $g$ in an empty droplet with cell barcode $b$. Upon sequencing, the UMI count for $g$ follows a Poisson distribution [1] with mean $\omega_b c_g \lambda_{bg}$, where $\omega_b$ represents the capture efficiency of library preparation for $b$ and $c_g$ represents the capture efficiency for transcripts of $g$. If we condition on the total count, the count vector for $b$ follows a multinomial distribution with probability of sampling $g$ equal to $c_g \lambda_{bg} C^{-1}$ where $C = \sum c_g \lambda_g$ across all $g$.

We further assume that $\lambda_{bg} \sim \text{Gamma}(v_b \mu_g, 1)$ independently for each $g$, where $\mu_g$ represents the ambient concentration of transcript molecules for $g$ and $v_b$ represents the volume of the droplet corresponding to $b$. Here, the absolute number of molecules is assumed to be sufficiently large such that $\lambda_{bg}$ is well-modelled by a continuous distribution. If the per-gene capture efficiency $c_g$ is similar across all $g$ and/or if the expected number of molecules $E(\lambda_{bg}) = v_b \mu_g$ is very large, the count vector for $b$ will follow a Dirichlet-multinomial distribution with $\alpha = v_b C_2$ and probability of sampling $g$ equal to $c_g \mu_g C_2^{-1}$ where $C_2 = \sum c_g \mu_g$. These conditions are necessary as scaling $\lambda_{bg}$ by $c_g$ does not preserve the Gamma scale parameter of 1 that is required to define a Dirichlet distrbution. If $c_g$ is constant across genes, it cancels out and can be ignored entirely, while if $v_b \mu_g$ is large, the variance of Dirichlet distribution approaches zero and any incorrect scaling of the variance due to $c_g$ can be ignored.

### 3 Efficient calculation of the Monte Carlo $p$-value

The naive approach to computing the Monte Carlo $p$-value for a barcode would be to directly sample $R$ count vectors from a Dirichlet-multinomial distribution, compute the likelihood $L'_{bi}$ for each count vector, and count the number of vectors with $L'_{bi} \leq L_b$. For $N$ genes, we would need $RN$ sampling operations to obtain the count vectors and the same number of multiplication operations to compute the likelihoods. For a dataset containing $B$ barcodes, this amounts to a total of $RNB$ operations.

When dealing with multiple barcodes, we can improve computational efficiency by using pre-existing results for barcodes with lower $t_b$. This is done by recognizing that sampling from a Dirichlet-multinomial distribution involves sampling from a multinomial distribution with a probability vector that itself is sampled from a Dirichlet distribution (i.e., each entry is sampled from a Gamma($\alpha\tilde{p}_g, 1$) distribution, and the sum of all entries is normalized to unity). Within a Monte Carlo iteration $i$:

1. We simulate an iteration-specific probability vector $\boldsymbol{\pi}_i = \{\pi_{1i}, \ldots, \pi_{gi}, \ldots, \pi_{Ni}\}$ across all genes from a Dirichlet distribution with a concentration parameter of $\alpha\tilde{p}_g$ for each gene $g$.

2. We consider the barcode $b_1$ with the lowest total count $t_{b_1}$. For this barcode, we sample a count vector $\mathbf{z}_i = \{z_{1i}, \ldots, z_{gi}, \ldots, z_{Ni}\}$ from a multinomial distribution with probabilities $\boldsymbol{\pi}_i$. We calculate the Dirichlet-multinomial likelihood $L'_{b_1 i}$ for $\mathbf{z}_i$ as previously described.

3. For barcode $b_2$ with the total $t_{b_2} = t_{b_1} + 1$, we sample a gene $k$ based on $\boldsymbol{\pi}_i$ and increment $z_{ki}$ to obtain an updated count vector. Similarly, we update the likelihood

$$L'_{b_2 i} = L'_{b_1 i} \left( \frac{t_{b_2}}{t_{b_2} + \alpha - 1} \right) \left( \frac{z_{ki} + \alpha\tilde{p}_k - 1}{z_{ki}} \right) ,$$

where $z_{ki}$ is the frequency of $k$ after the increment. (Iterative application of these updates can be used in cases where $t_{b_2} > t_{b_1} + 1$.) The updated $L'_{b_2 i}$ can be directly compared to $L_{b_2}$.

4. Repeat the previous step for all unique total counts in increasing order.

Repeating this procedure for $R$ iterations yields a distribution of likelihoods under the null hypothesis at each total count. This is used to obtain a Monte Carlo $p$-value for each barcode as previously described. The above approach only requires $R \max\{t_b\}$ sampling and multiplication operations, where each sampling step takes $\log(N)$ time due to a binary search across genes. It is independent of the number of barcodes and scales better than the naive approach with respect to the number of genes. Moreover, the maximum total count is usually low in droplet-based datasets (around 5000-10000). This results in a major reduction in computational work compared to the naive approach.

Our algorithm can be viewed as an extension of a simpler optimization where the set of likelihoods from all iterations are re-used for all barcodes with the same $t_b$. As a result, though, the Monte Carlo $p$-values are not strictly independent across barcodes. Instead, they are positively correlated as the same sampling choices are effectively shared between barcodes. This poses a theoretical problem for multiple testing correction with the Benjamini-Hochberg (BH) method, which relies on independent $p$-values. In practice, this is not a major issue as the BH method is robust to dependencies [2, 3]. Moreover, increasing $R$ reduces the imprecision of the $p$-values caused by random sampling. This means that the effect of correlated estimation errors between different $p$-values will become negligible at large $R$. We use $R = 10000$ by default, though this can be increased if the lower bound on the $p$-values [4] is greater than the threshold for significance after the Benjamini-Hochberg correction.

### 4 Detection differences between CellRanger and EmptyDrops

CellRanger also uniquely detects a number of barcodes with moderate total counts in some tested datasets, e.g., Supplementary Figure 10. This is attributable to the fact that these barcodes have expression profiles similar to the ambient pool, which prevents their detection by EmptyDrops. Without further information, it is difficult to determine whether the lack of detection is correct (i.e., the missing barcodes are empty droplets) or if this represents a minor loss of power. We note that EmptyDrops can always be made to report a superset of the barcodes identified by CellRanger, simply by setting $U$ to the CellRanger-identified threshold. We do not do so as the CellRanger threshold may not be appropriate. For example, CellRanger has an average observed FDR of 11 - 45% in simulations based on the T-cell dataset. By comparison, EmptyDrops approximately controls the type I error rate across assumed empty droplets in most real datasets (Supplementary Figure 1). Moreover, the CellRanger-only barcodes generally clustered with barcodes that were detected by both methods (Supplementary Figure 16a). This suggests that the "conservativeness" of EmptyDrops is not a major problem, as it only results in the loss of some cells from a cluster that would have been detected irrespectively.

**Supplementary Table 2:** Overview of the publicly available datasets from the 10X Genomics website that were used in this study. The organism, cell type, expected number of cells and the version of the CellRanger software (used for processing the raw sequencing data) is shown for each dataset.

| **Name** | **Organism** | **Cell type** | **Number** | **Version** |
|---|---|---|---:|---:|
| `pbmc4k` | Human | PBMCs | 4000 | 2.1.0 |
| `293t` | Human | 293T cell line | 2800 | 1.1.0 |
| `jurkat` | Human | Jurkat cell line | 3200 | 1.1.0 |
| `neuron_9k` | Mouse | Brain cells | 9000 | 2.1.0 |
| `neurons_900` | Mouse | Brain cells | 900 | 2.1.0 |
| `t_4k` | Human | Pan T cells | 4000 | 2.1.0 |

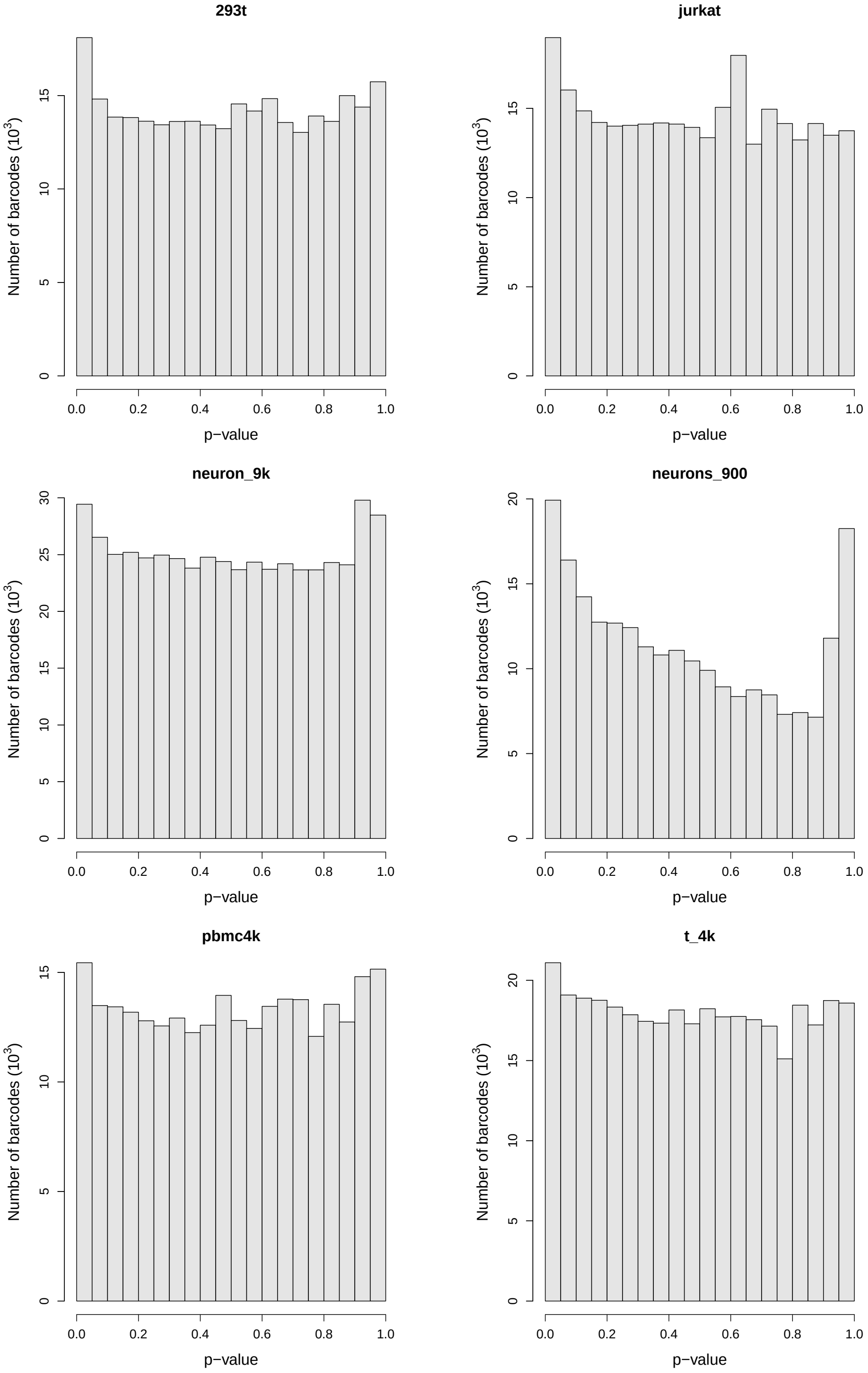


**Supplementary Figure 1:** Histograms of $p$-values for all barcodes with total UMI counts less than or equal to the threshold $T = 100$. Each plot corresponds to a dataset in Supplementary Table 2 where the $p$-value represents the deviation from the ambient profile.

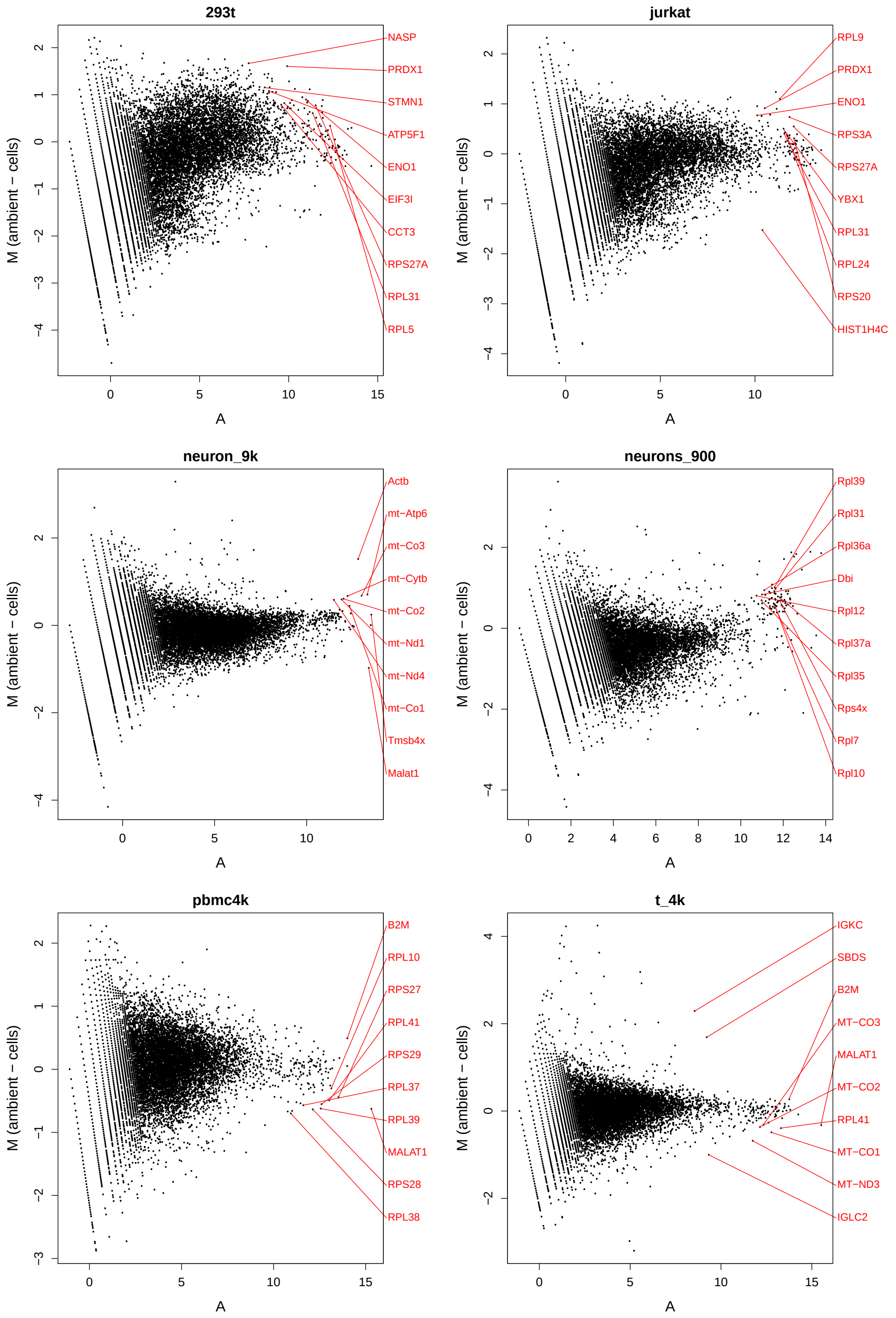


**Supplementary Figure 2:** MA plots for all genes comparing the mean expression profile of the ambient pool (all barcodes with total counts less than or equal to $T$) with the mean expression profile of all barcodes called as cells by EmptyDrops at an FDR of 0.1%. Each plot corresponds to a dataset in Supplementary Table 2. The top 10 genes with the strongest differential expression are also highlighted, based on a likelihood ratio test with a Poisson generalized linear model.

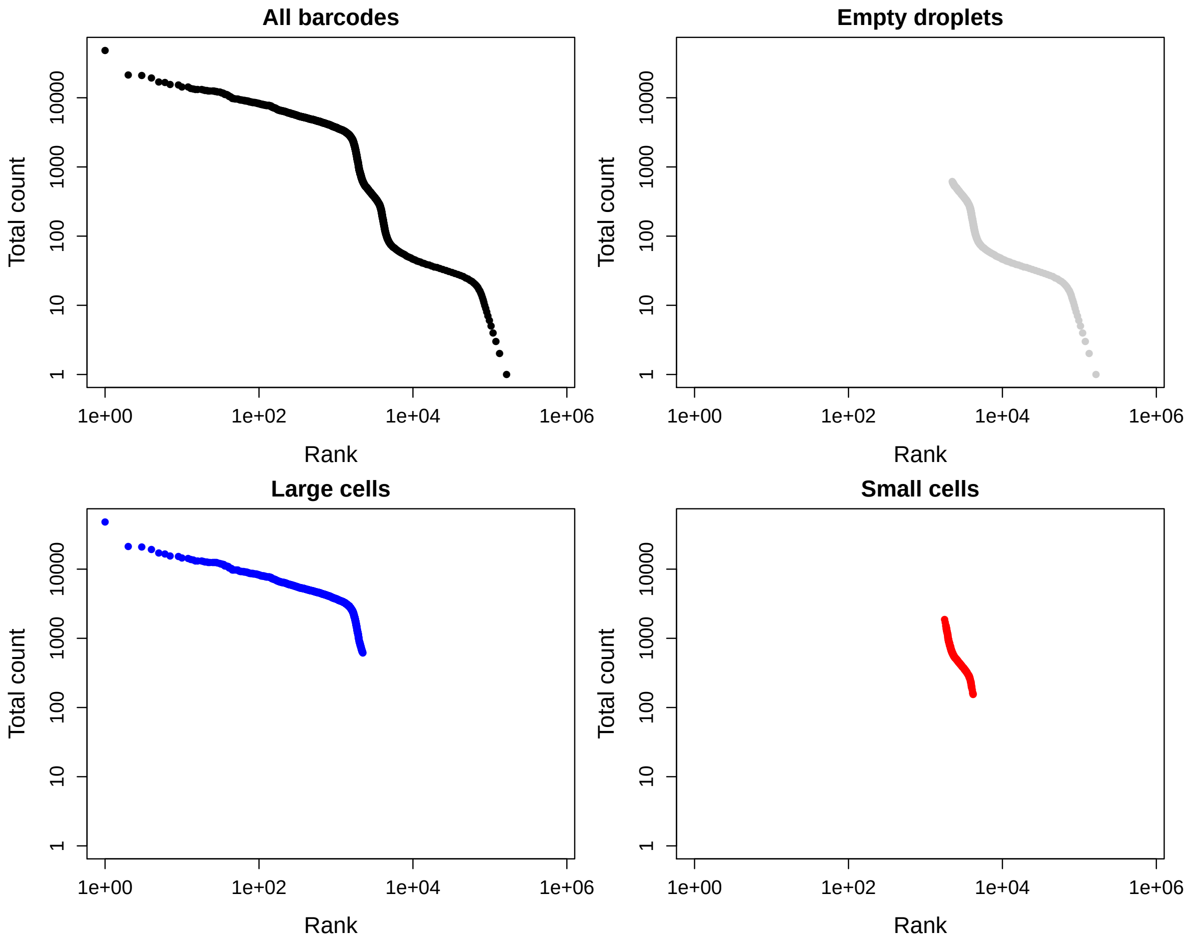


**Supplementary Figure 3:** Total count against the rank for each barcode in a simulation based on the PBMC dataset with $G_1 = G_2 = 2000$. Plots are shown for all barcodes, barcodes corresponding to empty droplets, and barcodes corresponding to large or small cells. Ranks are calculated from the entire set of barcodes in all plots, for ease of comparison between plots. All axes are on a log-scale.

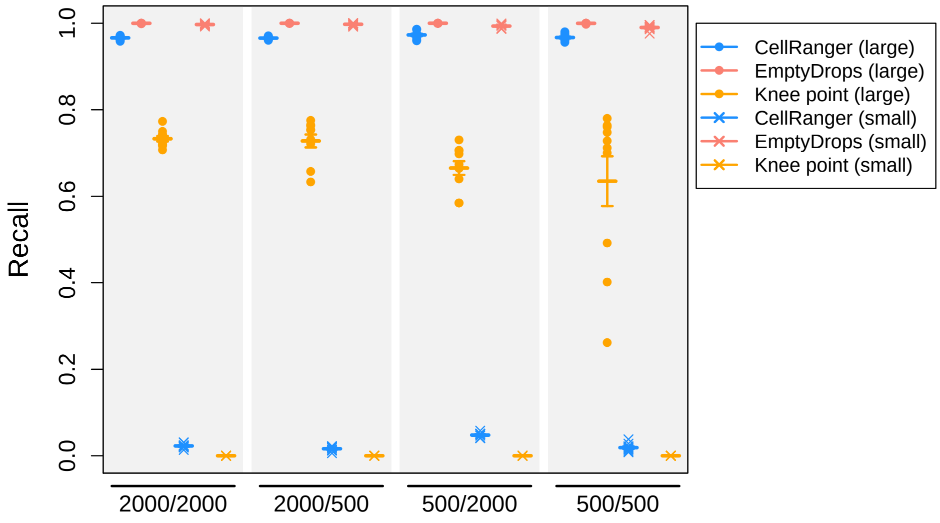


**(a)**

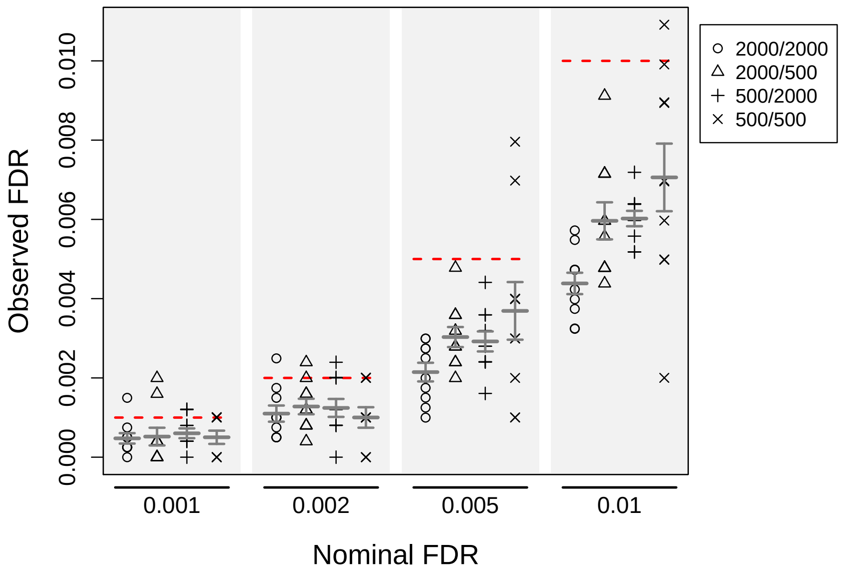


**(b)**

**Supplementary Figure 4:** Cell-calling results in simulations based on the 293T cell line dataset using either EmptyDrops or CellRanger. Simulation scenarios are labelled as $G_1/G_2$ where $G_1$ and $G_2$ are the number of barcodes in the group of large and small cells, respectively. (a) The recall for each method, defined as the proportion of detected cells from each group. EmptyDrops was used with an FDR of 0.1%. (b) The observed FDR in the set of libraries detected by EmptyDrops at a range of nominal FDR thresholds (dotted lines), defined as the proportion of detected droplets that are empty. In both plots, each point represents the result of one simulation iteration, the bar represents the mean value across 10 iterations and the error bars represent the standard error of the mean.

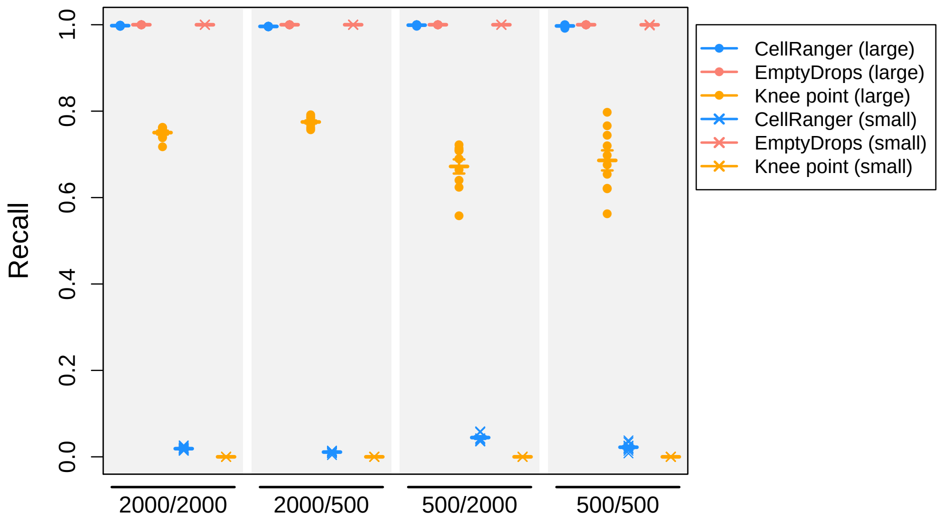


**(a)**

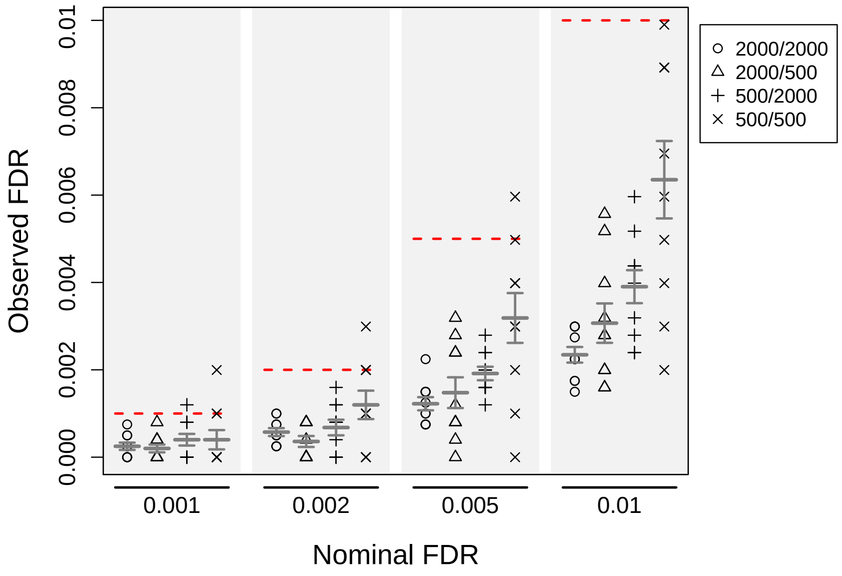


**(b)**

**Supplementary Figure 5:** Cell-calling results in simulations based on the Jurkat cell line dataset using either EmptyDrops or CellRanger. Simulation scenarios are labelled as $G_1/G_2$ where $G_1$ and $G_2$ are the number of barcodes in the group of large and small cells, respectively. (a) The recall for each method, defined as the proportion of detected cells from each group. EmptyDrops was used with an FDR of 0.1%. (b) The observed FDR in the set of libraries detected by EmptyDrops at a range of nominal FDR thresholds (dotted lines), defined as the proportion of detected droplets that are empty. In both plots, each point represents the result of one simulation iteration, the bar represents the mean value across 10 iterations and the error bars represent the standard error of the mean.

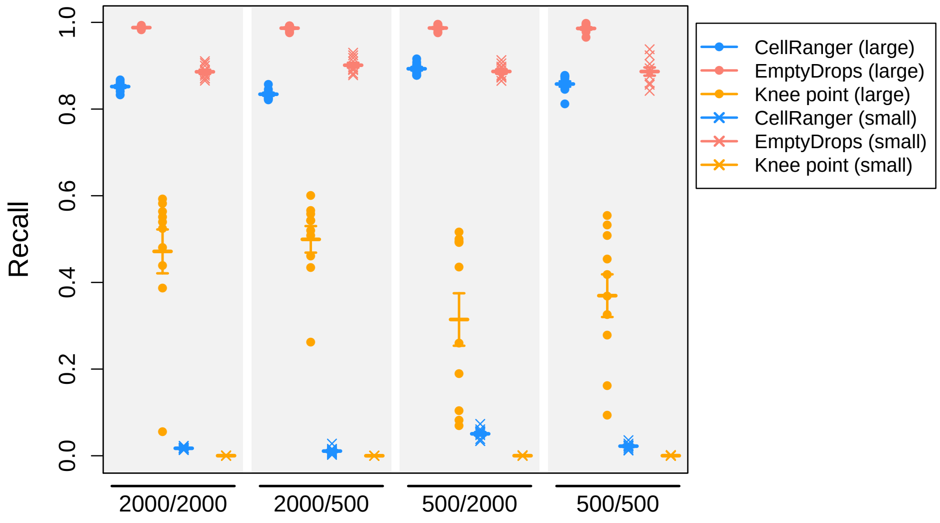


**(a)**

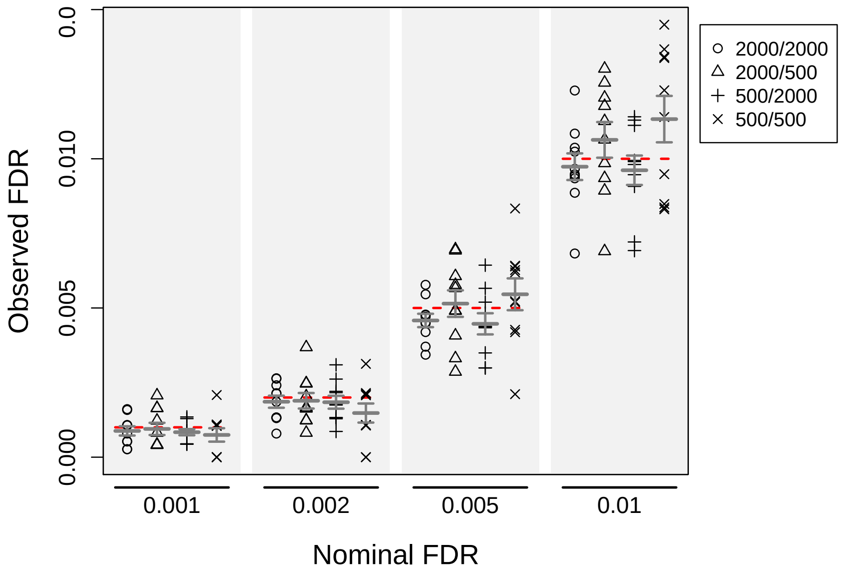


**(b)**

**Supplementary Figure 6:** Cell-calling results in simulations based on the 9K brain cell dataset using either EmptyDrops or CellRanger. Simulation scenarios are labelled as $G_1/G_2$ where $G_1$ and $G_2$ are the number of barcodes in the group of large and small cells, respectively. (a) The recall for each method, defined as the proportion of detected cells from each group. EmptyDrops was used with an FDR of 0.1%. (b) The observed FDR in the set of libraries detected by EmptyDrops at a range of nominal FDR thresholds (dotted lines), defined as the proportion of detected droplets that are empty. In both plots, each point represents the result of one simulation iteration, the bar represents the mean value across 10 iterations and the error bars represent the standard error of the mean.

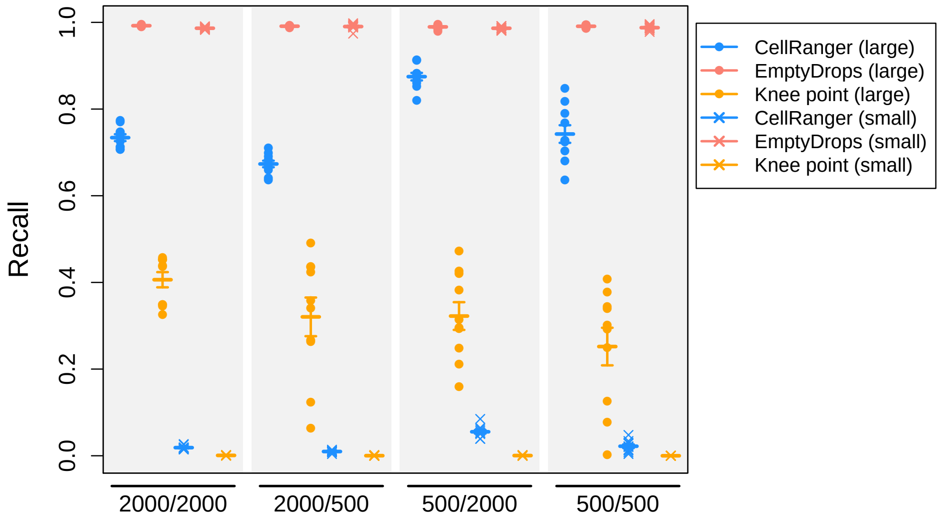


**(a)**

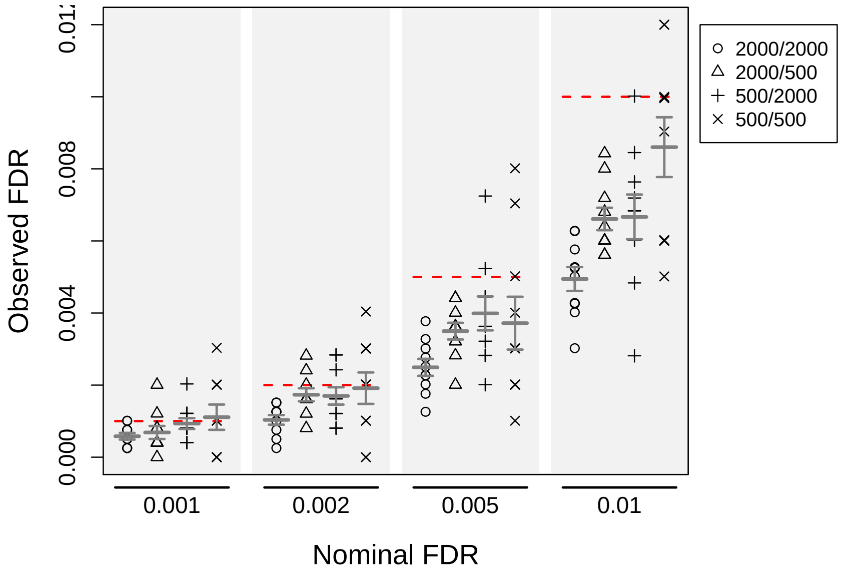


**(b)**

**Supplementary Figure 7:** Cell-calling results in simulations based on the 900 brain cell dataset using either EmptyDrops or CellRanger. Simulation scenarios are labelled as $G_1/G_2$ where $G_1$ and $G_2$ are the number of barcodes in the group of large and small cells, respectively. (a) The recall for each method, defined as the proportion of detected cells from each group. EmptyDrops was used with an FDR of 0.1%. (b) The observed FDR in the set of libraries detected by EmptyDrops at a range of nominal FDR thresholds (dotted lines), defined as the proportion of detected droplets that are empty. In both plots, each point represents the result of one simulation iteration, the bar represents the mean value across 10 iterations and the error bars represent the standard error of the mean.

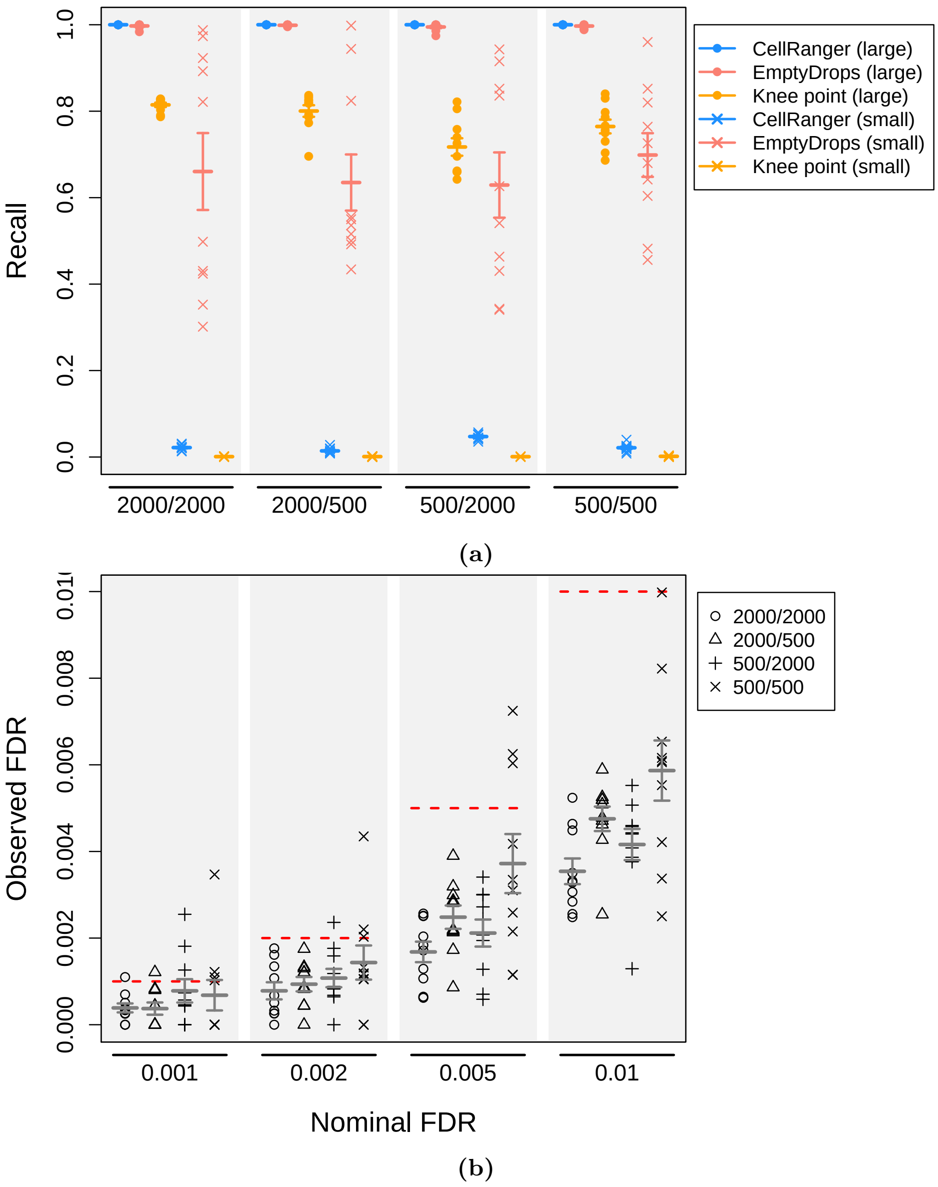


**Supplementary Figure 8:** Cell-calling results in simulations based on the pan T cell dataset using either EmptyDrops or CellRanger. Simulation scenarios are labelled as $G_1/G_2$ where $G_1$ and $G_2$ are the number of barcodes in the group of large and small cells, respectively. (a) The recall for each method, defined as the proportion of detected cells from each group. EmptyDrops was used with an FDR of 0.1%. (b) The observed FDR in the set of libraries detected by EmptyDrops at a range of nominal FDR thresholds (dotted lines), defined as the proportion of detected droplets that are empty. In both plots, each point represents the result of one simulation iteration, the bar represents the mean value across 10 iterations and the error bars represent the standard error of the mean.

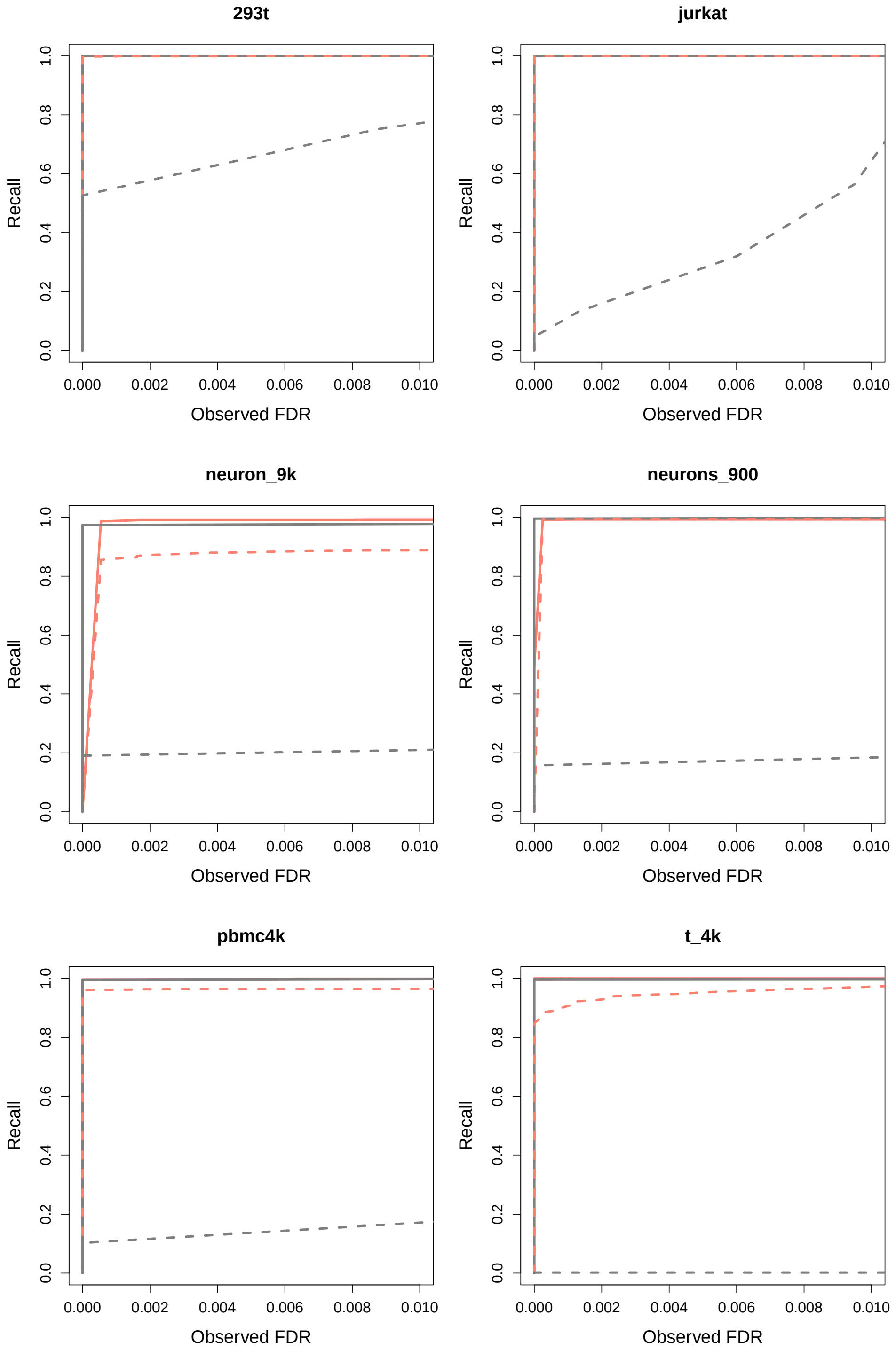


**Supplementary Figure 9:** Recall as a function of the observed FDR upon applying different cell calling strategies in simulations based on real datasets. For EmptyDrops (pink), the recall and observed FDR were computed at varying nominal FDR thresholds. For total count-based methods (grey), metrics were computed after increasing the total count threshold. Each simulation contained 2000 cells in both the large group (unbroken lines) and small group (dashed).

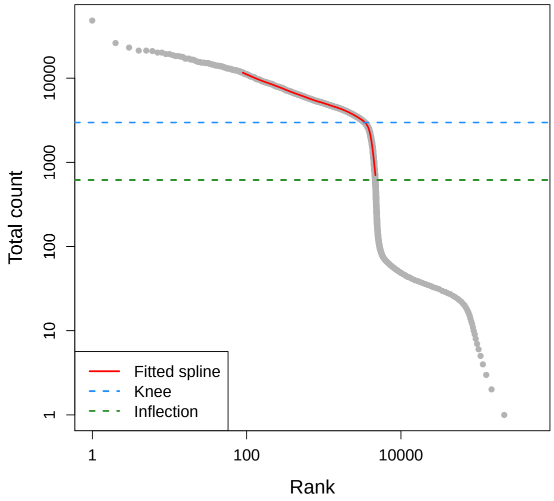


**(a)**

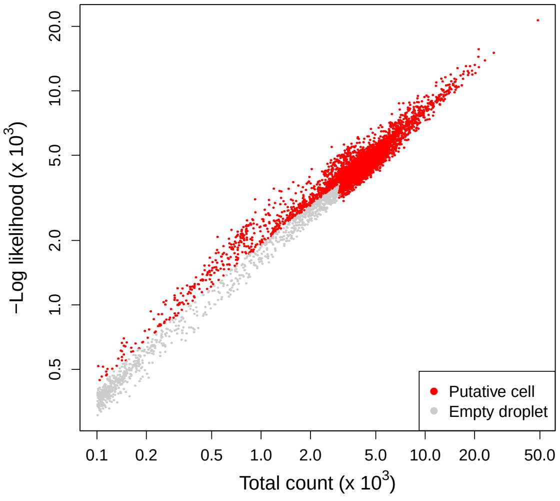


**(b)**

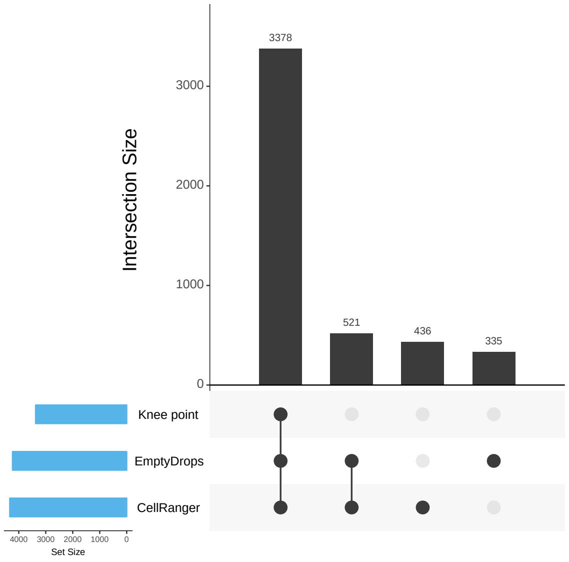


**(c)**

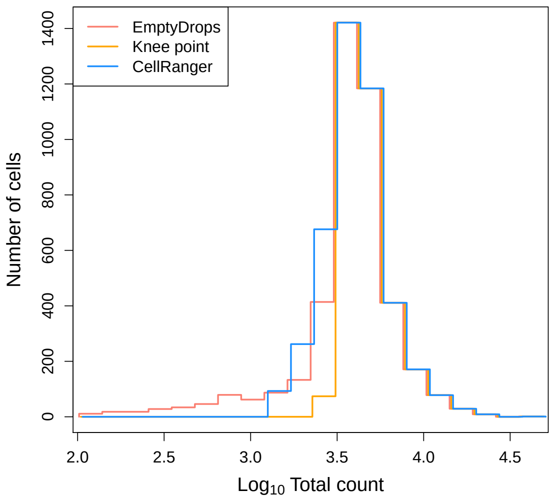


**(d)**

**Supplementary Figure 10:** Application of EmptyDrops and other cell detection methods to the 4K PBMC dataset. (a) A barcode rank plot showing the fitted spline used for knee point detection in EmptyDrops. The detected knee and inflection points are also shown. (b) The negative log-likelihood for each barcode in the multinomial model of EmptyDrops, plotted against the total count. Barcodes detected as putative cell-containing droplets at an FDR of 0.1% are labelled in red. Only barcodes with $t_b > T$ are shown. (c) An UpSet plot of the barcodes detected by each combination of methods (vertical bars). Horizontal bars represent the number of barcodes detected by each method. (d) Histogram outlines of the log-total count for barcodes detected by each method.

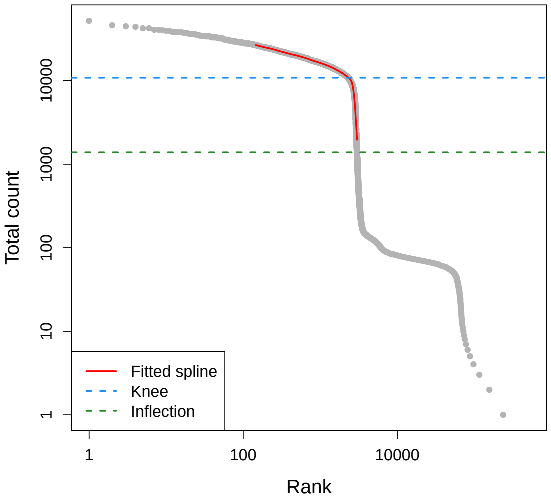


**(a)**

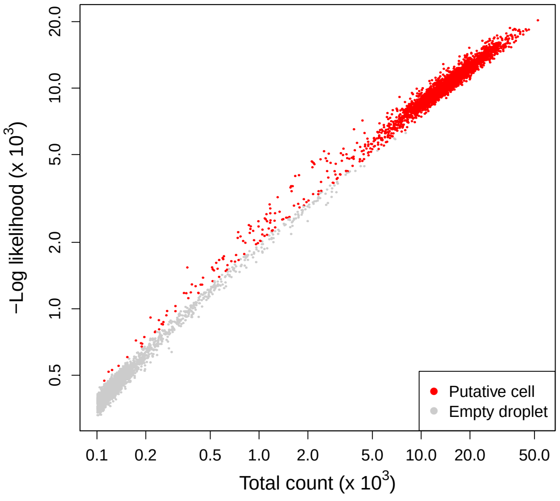


**(b)**

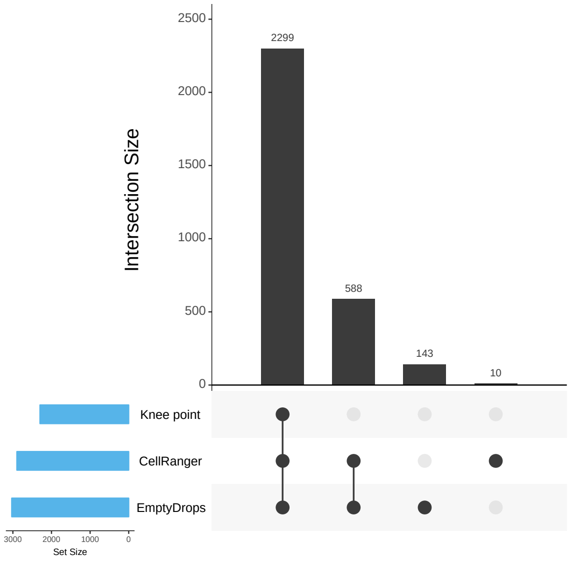


**(c)**

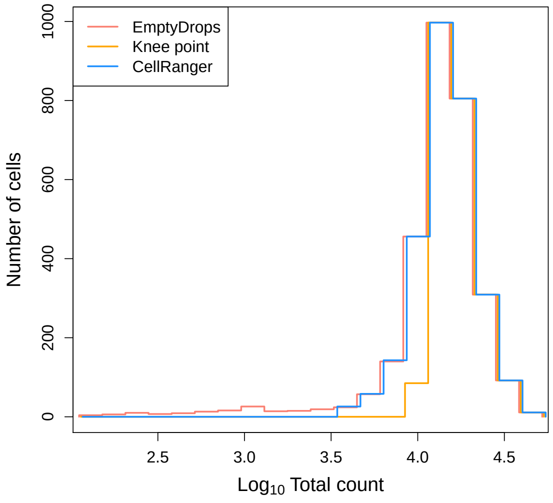


**(d)**

**Supplementary Figure 11:** Application of EmptyDrops and other cell detection methods to the 293T cell line dataset. (a) A barcode rank plot showing the fitted spline used for knee point detection in EmptyDrops. The detected knee and inflection points are also shown. (b) The negative log-likelihood for each barcode in the multinomial model of EmptyDrops, plotted against the total count. Barcodes detected as putative cell-containing droplets at an FDR of 0.1% are labelled in red. Only barcodes with $t_b > T$ are shown. (c) An UpSet plot of the barcodes detected by each combination of methods (vertical bars). Horizontal bars represent the number of barcodes detected by each method. (d) Histogram outlines of the log-total count for barcodes detected by each method.

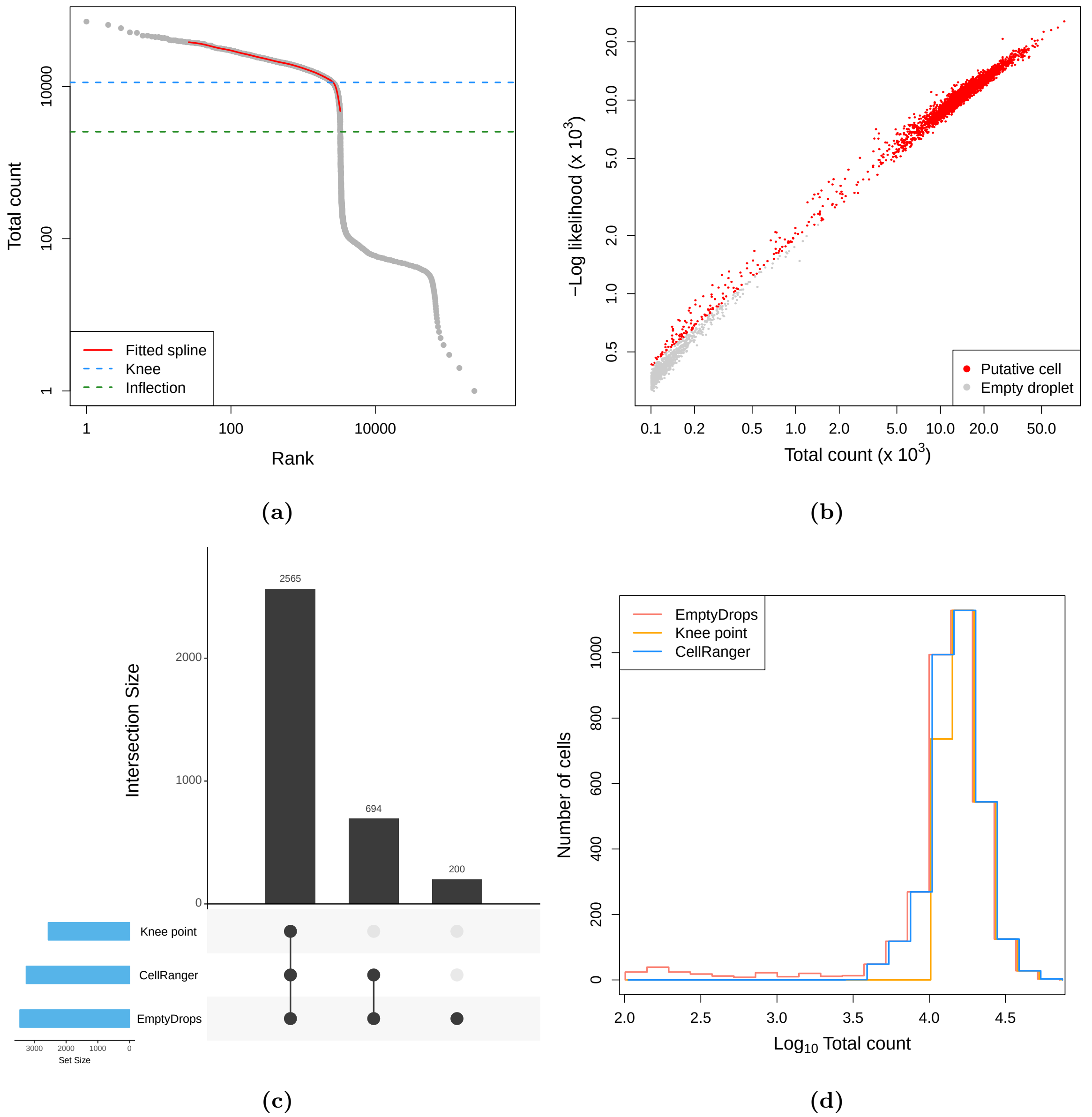


**Supplementary Figure 12:** Application of EmptyDrops and other cell detection methods to the Jurkat cell line dataset. (a) A barcode rank plot showing the fitted spline used for knee point detection in EmptyDrops. The detected knee and inflection points are also shown. (b) The negative log-likelihood for each barcode in the multinomial model of EmptyDrops, plotted against the total count. Barcodes detected as putative cell-containing droplets at an FDR of 0.1% are labelled in red. Only barcodes with $t_b > T$ are shown. (c) An UpSet plot of the barcodes detected by each combination of methods (vertical bars). Horizontal bars represent the number of barcodes detected by each method. (d) Histogram outlines of the log-total count for barcodes detected by each method.

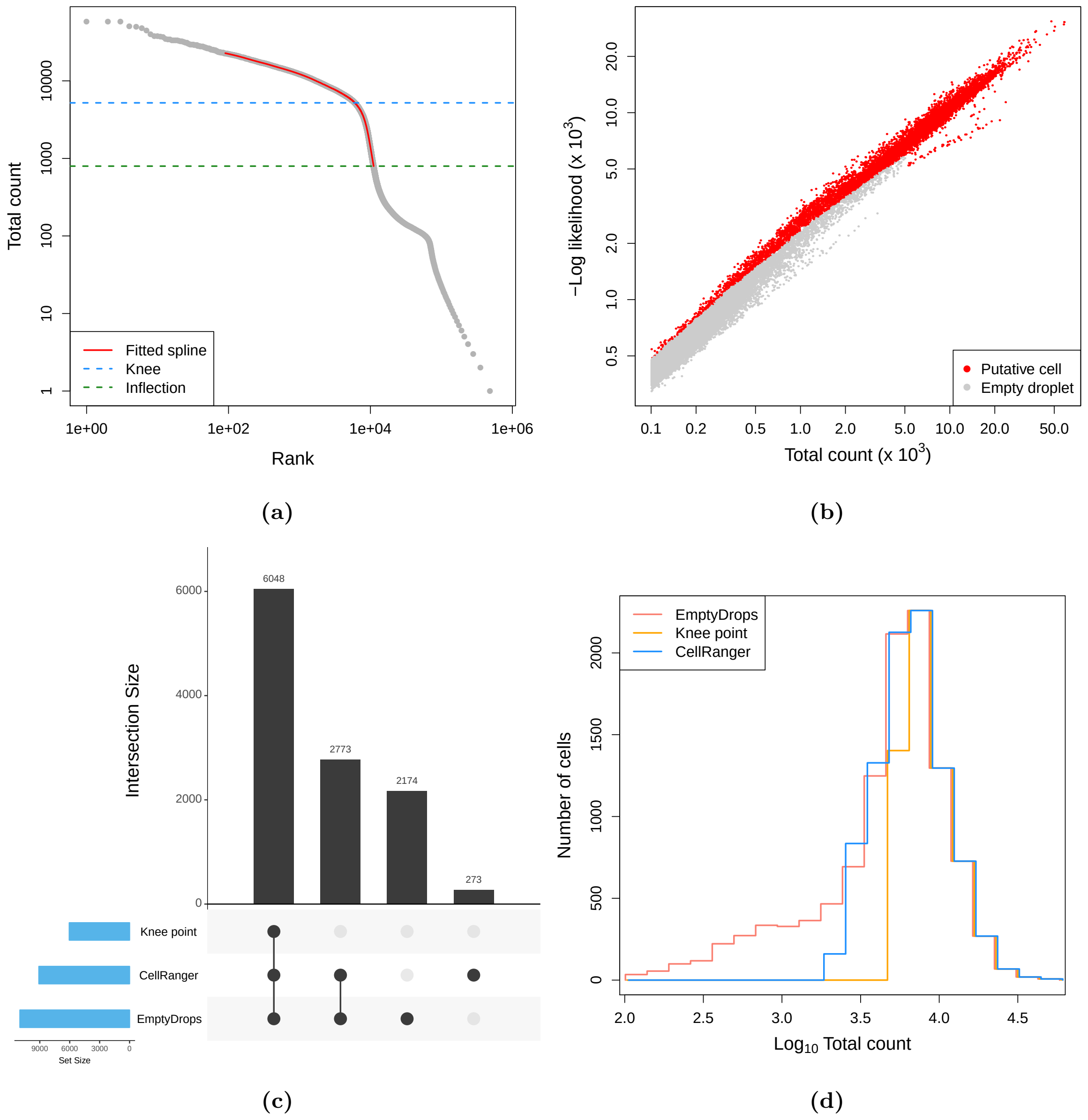


**Supplementary Figure 13:** Application of EmptyDrops and other cell detection methods to the 9K brain cell dataset. (a) A barcode rank plot showing the fitted spline used for knee point detection in EmptyDrops. The detected knee and inflection points are also shown. (b) The negative log-likelihood for each barcode in the multinomial model of EmptyDrops, plotted against the total count. Barcodes detected as putative cell-containing droplets at an FDR of 0.1% are labelled in red. Only barcodes with $t_b > T$ are shown. (c) An UpSet plot of the barcodes detected by each combination of methods (vertical bars). Horizontal bars represent the number of barcodes detected by each method. (d) Histogram outlines of the log-total count for barcodes detected by each method.

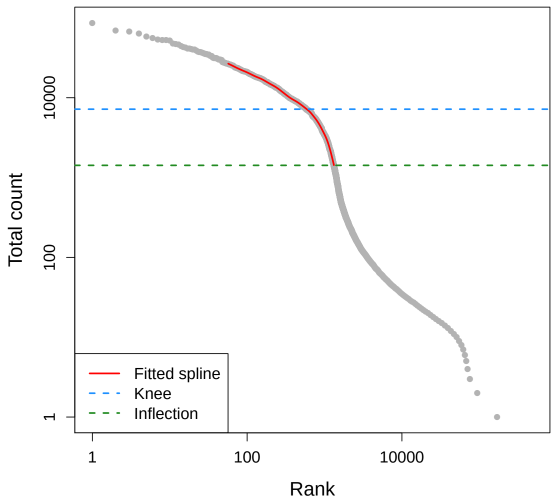


**(a)**

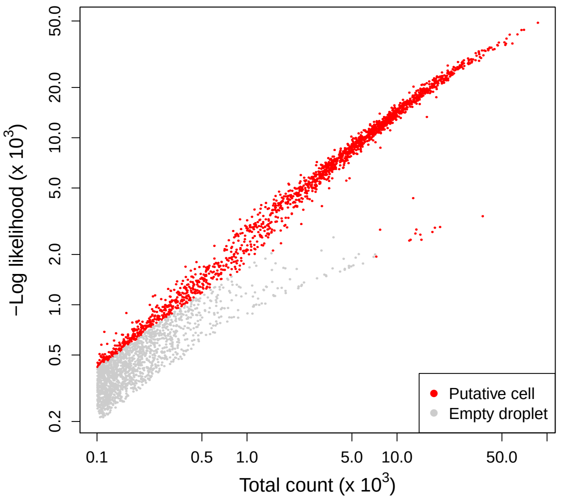


**(b)**

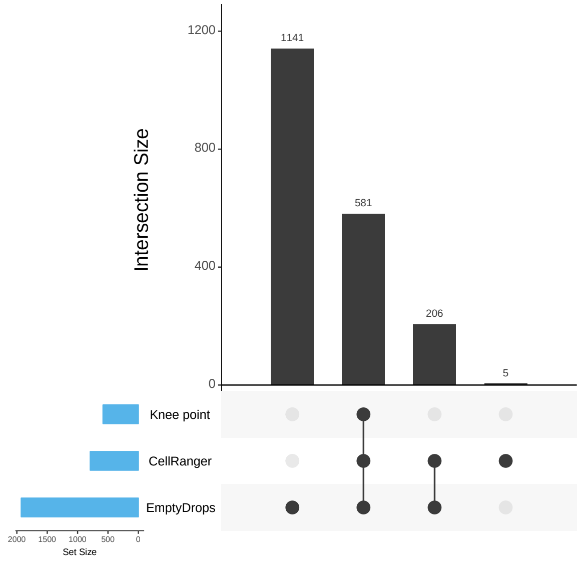


**(c)**

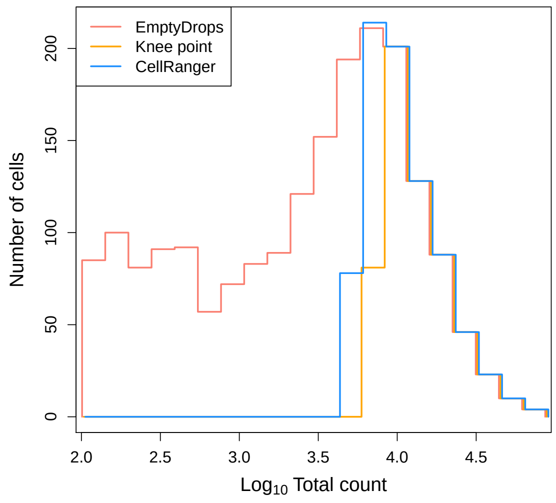


**(d)**

**Supplementary Figure 14:** Application of EmptyDrops and other cell detection methods to the 900 brain cell dataset. (a) A barcode rank plot showing the fitted spline used for knee point detection in EmptyDrops. The detected knee and inflection points are also shown. (b) The negative log-likelihood for each barcode in the multinomial model of EmptyDrops, plotted against the total count. Barcodes detected as putative cell-containing droplets at an FDR of 0.1% are labelled in red. Only barcodes with $t_b > T$ are shown. (c) An UpSet plot of the barcodes detected by each combination of methods (vertical bars). Horizontal bars represent the number of barcodes detected by each method. (d) Histogram outlines of the log-total count for barcodes detected by each method.

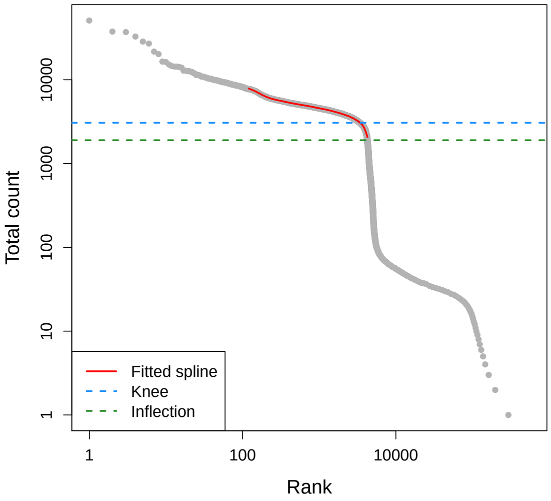


**(a)**

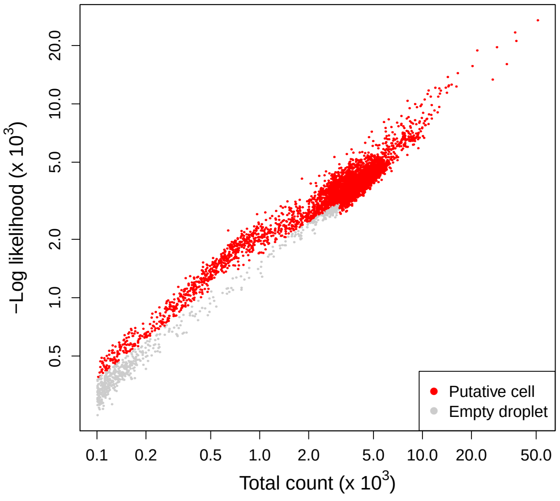


**(b)**

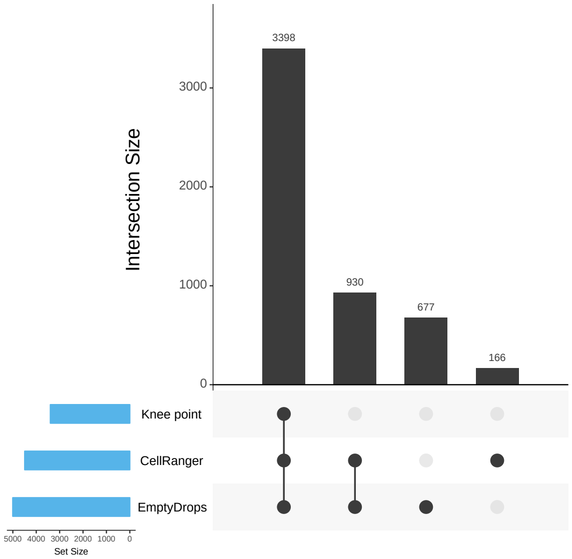


**(c)**

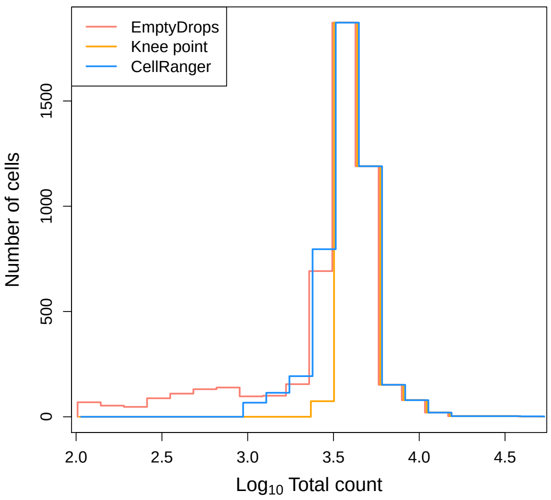


**(d)**

**Supplementary Figure 15:** Application of EmptyDrops and other cell detection methods to the pan T cell dataset. (a) A barcode rank plot showing the fitted spline used for knee point detection in EmptyDrops. The detected knee and inflection points are also shown. (b) The negative log-likelihood for each barcode in the multinomial model of EmptyDrops, plotted against the total count. Barcodes detected as putative cell-containing droplets at an FDR of 0.1% are labelled in red. Only barcodes with $t_b > T$ are shown. (c) An UpSet plot of the barcodes detected by each combination of methods (vertical bars). Horizontal bars represent the number of barcodes detected by each method. (d) Histogram outlines of the log-total count for barcodes detected by each method.

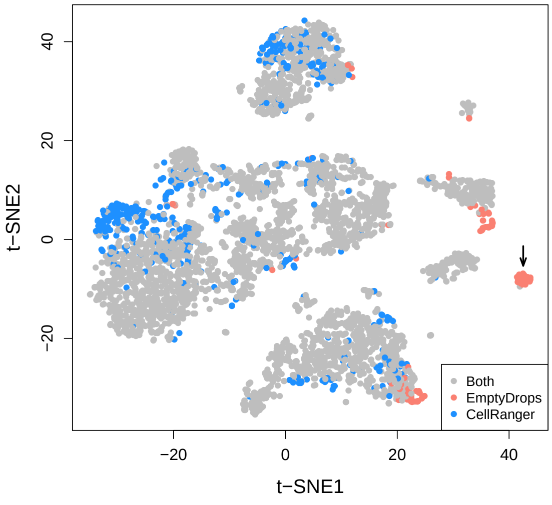


**(a)**

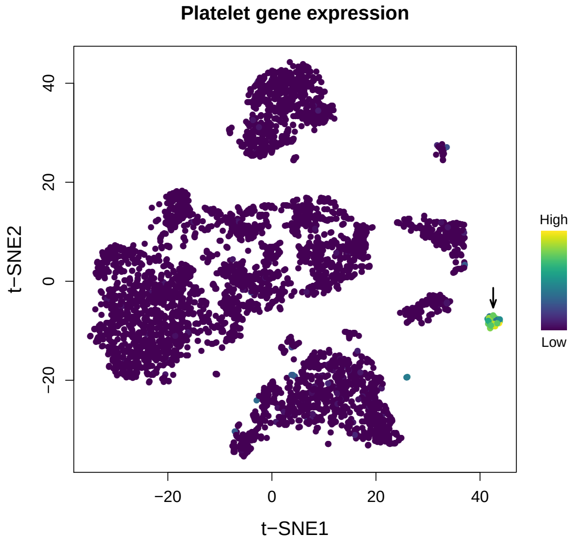


**(b)**

**Supplementary Figure 16:** *t*-SNE plots for the 4K PBMC dataset, constructed from barcodes that were detected with EmptyDrops and/or CellRanger. Each point represents a barcode and is coloured based on (a) whether it was detected as a cell with each method; or (b) the expression of platelet marker genes *PF4* and *PPBP*. Expression in each barcode was quantified as the sum of the normalized log-expression values across all marker genes. Arrows mark the putative platelet population.
